# Age-dependent contribution of domain-general networks to semantic cognition

**DOI:** 10.1101/2020.11.06.371153

**Authors:** Sandra Martin, Dorothee Saur, Gesa Hartwigsen

## Abstract

Aging is characterized by a decline of cognitive control. In semantic cognition, this leads to the paradox that older adults usually show poorer task performance than young adults despite their greater semantic knowledge. So far, the underlying neural changes of these behavioral differences are poorly understood. In the current neuroimaging study, we investigated the contribution of domain-general networks to a verbal semantic fluency task in young and older adults. In both age groups, task processing was characterized by a strong positive integration within the multiple-demand as well as between the multiple-demand and the default mode system during semantic fluency. However, the behavioral relevance of strengthened connectivity differed between groups: While within-network functional connectivity in both networks predicted greater efficiency in semantic fluency in young adults, it was associated with a slower performance in older adults. Moreover, only young adults profited from increased connectivity between networks for their semantic memory performance. Our results suggest that the functional coupling of usually anti-correlated networks is critical for successful task processing, independent of age, when access to semantic memory is required. Furthermore, our findings lend novel support to the notion of reduced efficiency in the aging brain, due to neural dedifferentiation in semantic cognition.

## Introduction

Semantic cognition is a fundamental human ability that is central to communication across the life span. Key facets of semantic cognition refer to our knowledge of the world and the meaning of words and sentences. With respect to cognitive changes across the adult life span, cognitive control processes – also referred to as fluid intelligence – are well established to steadily decline with increasing age (Hedden and Gabrieli, 2004) whereas semantic knowledge (so-called crystallized intelligence) has been shown to remain stable or might even increase with age due to the ongoing accrual of knowledge and experience across the life course (Verhaegen, 2003). In the domain of semantic cognition, the impact of aging thus seems to depend on both the specific cognitive demand of a task and the individual semantic knowledge.

At the neural level, cognitive changes with age are mirrored by large-scale reorganization processes at the structural and functional levels (Grady, 2012; Morcom and Johnson, 2015). Task-related performance changes in older adults have been associated with a pattern of dedifferentiation of neural activity (Park et al., 2004; Li et al., 2001) which is reflected by an under-recruitment of domain-specific regions (Lövdén et al., 2010) and reduced task-specific lateralization (Cabeza, 2002). Dedifferentiation is further characterized by an increased recruitment of areas in the domain-general multiple-demand network (MDN; Lövdén et al., 2010) and a reduced deactivation of regions in the default mode network (DMN; Andrews-Hanna et al., 2007; Damoiseaux et al., 2008; Persson et al., 2007). A recent meta-analysis that investigated age-related effects on the neural substrates of semantic cognition confirmed the upregulation of the MDN in older adults for a variety of semantic tasks (Hoffman and Morcom, 2018).

In addition to local changes in task-related activity, alterations in the functional connectivity of large-scale neural networks have become a hallmark of brain aging (Damoiseaux et al., 2017; Li et al., 2015; Spreng et al., 2016). A common observation is that functional network segregation declines with age which is evident in the form of decreased within- and increased between-network functional connectivity (Chan et al., 2014; Geerligs et al., 2015; Spreng et al., 2016). These changes have been interpreted as dedifferentiation of network interactions, paralleling local task-related neural changes (Spreng and Turner, 2019). However, the majority of studies investigated these changes at rest and it is thus less clear how aging affects task-related functional connectivity and how this is associated with behavior.

The recently proposed default-executive coupling hypothesis of aging (DECHA; Turner and Spreng, 2015; Spreng and Turner, 2019) suggests that the observed activity increase in MDN regions and the reduced deactivation of the DMN co-occur and are functionally coupled in older adults. This shift in network coupling is based on the accrual of semantic knowledge and the parallel decline of cognitive control abilities. Older adults thus rely more strongly on their preserved semantic knowledge which is reflected by a reduced deactivation of DMN regions compared to young adults. According to this hypothesis, context and cognitive demand of a task determine if this increased default-executive coupling in older adults is beneficial or maladaptive. On this basis, the framework predicts a stable performance in older adults for tasks that rely on crystallized intelligence in the form of intrinsic prior knowledge and require little cognitive control whereas externally-directed cognitive tasks result in poorer performance in older adults due to their dependence on control resources.

So far, the integration of domain-general networks in semantic word retrieval in older adulthood is poorly understood. In this context, semantic fluency tasks, which require participants to generate words that belong to a specific category within a given time, provide a unique opportunity since they require an interaction of verbal semantic and general cognitive control processes (Gordon et al., 2018; Schmidt et al., 2017; Whiteside et al., 2016). Semantic fluency tasks test a natural and important communicative ability as they rely on accessing related concepts to retrieve words. Furthermore, semantic fluency is of high ecological validity, for example when writing a shopping list (Shao et al., 2014), and is frequently implemented as a measure of language and neuropsychological abilities in healthy as well as clinical populations (Schmidt et al., 2017). The impact of aging on semantic fluency is especially interesting since its strong link to semantic memory would predict preserved performance for older adults. Yet, the opposite pattern is usually observed, suggesting a high load on cognitive control processes for this task (e.g., Kavé and Knafo-Noam, 2015; Gordon et al., 2018; Troyer et al., 1997). Most studies that implemented semantic fluency tasks in neuroimaging experiments reported age-related changes within domain-specific networks (Baciu et al., 2016; Marsolais et al., 2014) or pre-defined regions of interest, mainly in the prefrontal cortex (Meinzer et al., 2009; Meinzer et al., 2012a; Meinzer et al., 2012b). However, based on the outlined changes in semantic and cognitive control abilities, older adults could show a shift in network coupling with a stronger engagement of domain-general networks which might be additionally modulated by task demand.

Thus, the aim of the present study was to explore and compare the interplay of domain-specific and domain-general networks during word retrieval in healthy young and older adults. We implemented an fMRI study with a paced overt semantic fluency task, which included an explicit modulation of task difficulty. A counting task was used as a low-level verbal control task. First, we were interested in delineating the network for semantic fluency and its interaction with task demand. Secondly, we asked whether age modulates both activation patterns and behavioral performance. Finally, we were interested in task-related functional interactions between domain-specific and domain-general networks. To this end, we combined univariate whole-brain analyses with generalized psycho-physiological interaction (gPPI) analyses. We applied traditional gPPI analyses to explore the functional coupling of the strongest activation peaks for semantic fluency. This allowed us to investigate the age-related contribution of domain-general networks to language processing. Furthermore, we used modified gPPI analyses to examine functional connectivity within and between regions of domain-general networks. We were interested in age-related effects on functional connectivity patterns and how within- and between-network functional connectivity relate to behavioral performance for both age groups. We expected a language-specific left-lateralized network for semantic fluency. We further hypothesized that increased task demand (reflected by the contrast of semantic fluency with counting, as well as by the modulation of difficulty within the semantic fluency task) would affect task performance and should be accompanied by an increased recruitment of domain-general systems. With respect to task-related functional connectivity, we reasoned that older adults should demonstrate a stronger involvement of the DMN for the semantic fluency task based on their increased semantic knowledge. Moreover, a higher task load associated with general cognitive decline might further result in a stronger recruitment of the MDN in older adults. However, in line with neurocognitive theories of aging, the increased recruitment of domain-general systems might be associated with reduced specificity and efficiency; thus, overall leading to a poorer performance in the older adults.

## Materials and Methods

### Participants

Twenty-eight healthy older adults (mean age: 65.2 years, range: 60-69 years) and thirty healthy younger adults (mean age: 27.6 years, range: 21-34 years) completed our study. The data of 3 additional participants in the older group as well as individual runs of 6 participants were excluded due to excessive head movement during fMRI (> 1 voxel size). Groups were matched for gender. Participants in the younger group had significantly more years of education (*t*(55.86) = 5.2, *p* < .001). All participants were native German speakers and right-handed according to the Edinburgh Handedness Inventory (Oldfield, 1971). They had normal hearing, normal or corrected-to-normal vision, and no history of neurological or psychiatric conditions or contraindication to MR-scanning. Older adults were additionally screened for cognitive impairments using the Mini-Mental State Examination (Folstein et al., 1975; all ≥26/30 points) and for depression with the Beck Depression Inventory (Beck et al., 1996; all ≤14/ points). The study was approved by the local ethics committee of the University of Leipzig and conducted in accordance with the Declaration of Helsinki. Participants gave written informed consent prior to the experiment. They received 10 Euro per hour for their participation.

### Neuropsychological Assessment

A battery of neuropsychological tests was administered to all participants to assess cognitive functioning. Verbal knowledge and executive language functions were measured with the German version of the spot-the-word test (Wortschatztest; Schmidt and Metzler, 1992; Baddeley et al., 1993), a German version of the reading span test (Daneman and Carpenter, 1980), and the semantic subtest of a verbal fluency test (Regensburger Wortflüssigkeitstest; Aschenbrenner et al., 2000). The latter comprised two 1-minute trials of semantic categories (surnames and hobbies) that were not part of the fMRI task. Additionally, executive functions were assessed with the Digit Symbol Substitution Test (Wechsler, 1944) and the Trail Making Test A/B (Reitan, 1958).

Group comparisons showed that older participants only performed better than the younger group on the spot-the-word test (Table 1 and Supplementary Fig. S1), which is considered a measure of lexical semantic knowledge and vocabulary. Consistent with our results, it has been shown to be robust to aging and cognitive decline (Baddeley et al., 1993; Law and O’Carroll, 1998; Cohen-Shikora and Balota, 2016). Our results confirm the maintenance of semantic memory across age (Grady, 2012) and an increase in size of vocabulary with age (Verhaegen, 2003). All other tests showed better performance for younger participants, which is in line with the assumption of a general decline in executive functions like working memory and processing speed with age (e.g., Balota et al., 2000; Zacks et al., 2000). However, when considering age-corrected norms, the older participants performed within normal ranges on all neuropsychological tests.

**Table 1.** Demographic and neuropsychological characteristics of participants

|  | Young adults<br>(*n* = 30) | Older adults<br>(*n* = 28) |
| --- | --- | --- |
| *Demographics* |  |  |
| Age (years) | 27.6 (4.4) | 65.2 (2.8) |
| Gender (F:M) | 16:14 | 14:14 |
| Education (years) | 18.7 (2.6) | 15.2 (2.5)* |
| Beck Depression Inventory (cut-off 18 points) | – | 4.7 (4.1) |
| *Neuropsychological* |  |  |
| Spot-the-word test (max. 40) | 29.1 (3.2) | 31.5 (2.5)* |
| Semantic fluency (sum surnames, hobbies) | 51.2 (8.4) | 40.7 (6.7)* |
| Reading span test (max. 6) | 3.5 (1) | 2.9 (0.7)* |
| Digit symbol substitution test (max. 90 in 90 s) | 72.1 (11.4) | 50.2 (10.4)* |
| Trail Making Test A (time in s) | 17.3 (5.8) | 25.4 (6.4)* |
| Trail Making Test B (time in s) | 36.1 (11.9) | 61.8 (29.4)* |
| Mini-Mental State Examination (max. 30 points) | – | 28.36 (1.2) |

Note: Mean values of raw scores with standard deviations. \* Significant differences between age groups at $p < 0.01$ .

### Experimental Design

All participants completed one fMRI session which was divided into two runs. Tasks consisted of a paced overt semantic fluency task and a control task of paced counting which were implemented in a block design in the scanner. We chose a paced design for our tasks since it has been shown to be less sensitive to motion artifacts and to yield robust brain activation patterns (Basho et al., 2007). Task blocks were 43 s long and separated by rest blocks of 16 s (Fig. 1A). Each block started with a 2 s visual word cue indicating whether participants were expected to generate category exemplars or count forward (1 to 9) or backward (9 to 1). This was followed by 9 consecutive trials of the same category or counting task, respectively. Trials within one block were separated by inter-stimulus intervals of 2-4 s. Participants were instructed to generate one exemplar for a category or one number per trial, which was indicated through a green cross on the screen, and to pause when the cross turned red (Fig. 1B and C). They were told not to repeat items and to say “next” if they could not think of an exemplar for the respective category. Each run contained 10 semantic fluency blocks, divided in easy and difficult categories, and 10 counting blocks, consisting of forward and backward counting, thus resulting in a total duration of 19.4 min per run. The order of blocks was counter-balanced and pseudo-randomized across participants. Before the fMRI experiment, participants received instructions and practiced the task with a separate set of categories outside the scanner. Stimuli were presented using the software Presentation (Neurobehavioral Systems, Berkeley, USA; version 18.0). Answers were recorded via a FOMRI III microphone (Optoacoustics, Yehuda, Israel).

**Figure 1.**
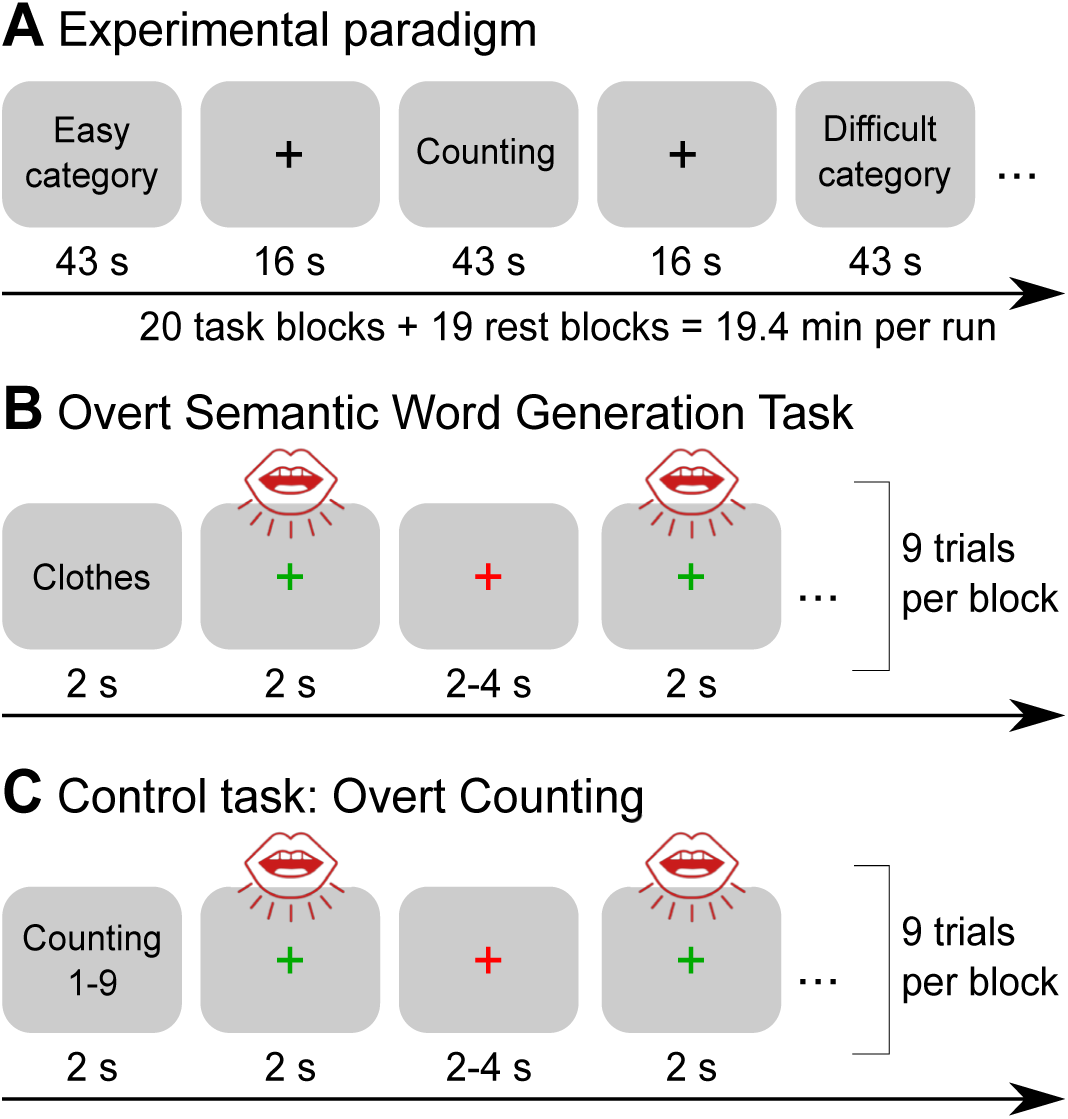
Experimental design. (A) fMRI experiment consisting of alternating blocks of a semantic fluency and a counting task separated by 16-s rest periods. (B) and (C) demonstrate the implementation of the paced design in both tasks. Procedures were identical for both tasks. Participants were instructed to produce one exemplar for a category or to say one number per green cross, respectively, and to pause when the cross turned red. Each block contained 9 trials which were separated by jittered inter-stimulus intervals.

### Stimuli

Stimuli consisted of 20 semantic categories which were divided in 10 easy and 10 difficult categories. Difficulty was assessed in a separate pilot study with 24 older adults (10 male, mean age: 65 years, range: 60-69 years). Participants were recruited and screened using similar criteria as in the fMRI study. They generated as many exemplars as possible during 1-minute trials for 30 semantic categories which were taken from German category-production norm studies (Mannhaupt, 1983; Glauer et al., 2007). Responses were recorded and subsequently transcribed and analyzed. Based on the total number of correct exemplars produced for each category, the 10 categories with the largest number of produced items (colors, body parts, clothing, types of sport, animals, car parts, professions, trees, food, and musical instruments) and the 10 categories with the fewest items (flowers, insects, metals, kitchen devices, tools, gardening tools, fishes, cosmetics, toys, and sweets) were chosen for the easy and difficult conditions of the semantic fluency task in the fMRI experiment, respectively. Easy (*M* = 16.8, *SD* = 2.5) and difficult categories (*M* = 10.4, *SD* = 2) differed significantly in the mean number of generated exemplars (*t*(38.9) = 9.2, *p* < 0.001) during piloting.

### Data Acquisition and Preprocessing

MR images were collected at a 3-Tesla Prisma Scanner (Siemens, Erlangen, Germany) with a 32-channel head coil. For the acquisition of fMRI data, a dual gradient echo-planar imaging multiband sequence (Feinberg et al., 2010) was used for optimal BOLD sensitivity across the whole brain (Poser et al., 2006; Halai et al., 2014). The following scanning parameters were applied: TR = 2,000 ms; TE = 12 ms, 33 ms; flip angle = 90°; voxel size = 2.5 x 2.5 x 2.5 mm with an inter-slice gap of 0.25 mm; FOV = 204 mm; multiband acceleration factor = 2. To increase coverage of anterior temporal lobe (ATL) regions, slices were tilted by 10° of the AC-PC line. 616 images consisting of 60 axial slices in interleaved order covering the whole brain were continuously acquired per run. Additionally, field maps were obtained for later distortion correction (TR = 8000 ms; TE = 50 ms). This study analyzed the data from echo 2 (TE = 33 ms) since preprocessing was performed using the software fMRIPrep (Esteban et al., 2019), which currently does not support the combination of images acquired at different echo times. We chose to use results from preprocessing with fMRIPrep since this pipeline provides state-of-the-art data processing while allowing for full transparency and reproducibility of the applied methods and a comprehensive quality assessment of each processing step which facilitates the identification of potential outliers. We also double-checked results from preprocessing with fMRIPrep with a conventional SPM preprocessing pipeline of both echoes. The comparison of both pipelines did not reveal big differences in analysis results. A high-resolution, T1-weighted 3D volume was obtained from our in-house database (if it was not older than 2 years) or collected after the functional scans using an MPRAGE sequence (176 slices in sagittal orientation; TR = 2300 ms; TE = 2.98 ms; flip angle = 9°; voxel size = 1 x 1 x 1 mm; no slice gap; FOV = 256 mm).

Preprocessing was performed using fMRIPprep 1.2.6 (Esteban et al., 2019), which is based on Nipype 1.1.7 (Gorgolewski et al., 2017). In short, within the pipeline, anatomical images were processed using the software ANTs (Tustison et al., 2010) for bias field correction, skull stripping, coregistration, and normalization to the skull-stripped ICBM 152 Nonlinear Asymmetrical template version 2009c (Fonov et al. 2009). FreeSurfer (Dale, Fischl, and Sereno 1999) was used for brain surface reconstruction and FSL (Jenkinson et al. 2012) for segmentation. Functional data of each run were skull stripped, distortion corrected, slice-time corrected, coregistered to the corresponding T1 weighted volume, and resampled to MNI152NLin2009cAsym standard space. Head-motion parameters with respect to the BOLD reference (transformation matrices, and six corresponding rotation and translation parameters) were estimated before any spatiotemporal filtering using FSL. For more details of the pipeline, see the section corresponding to workflows in fMRIPrep’s documentation (https://fmriprep.org/en/1.2.6/workflows.html). After preprocessing, 29 volumes from the beginning of each run were discarded since they were collected for the combination of the short and long TE images via an estimation of the temporal signal-to-noise-ratio (Poser et al., 2006). This yielded 587 normalized images per run which were included in further analyses. The images were smoothed with a 5 mm^3^ FWHM Gaussian kernel using Statistical Parametrical Mapping software (SPM12; Wellcome Trust Centre for Neuroimaging), implemented in MATLAB (version 9.3/2017b).

### Data Analysis

#### Behavioral data

Response recordings during the semantic fluency task were cleaned from scanner noise using Audacity (version 2.3.2, https://www.audacityteam.org/) and verbal answers and onset times were transcribed by 3 independent raters. Repetitions of words within a category were counted as incorrect, incomplete answers and null reactions were marked separately, and full categories that had been missed by participants (in total 10 categories) were excluded from the analyses. Statistical analyses were performed with R via RStudio (R Core Team, 2018) and the packages lme4 (Bates et al., 2015) for mixed models and ggplot2 (Wickham, 2016) for visualizations. We used sum coding (ANOVA-style encoding) for all categorical predictors. In this way, the intercept represents the mean across conditions (grand mean), and the model coefficients represent the difference between the grand mean and the mean of the respective condition. For the analysis of accuracy, a generalized linear mixed-effects logistic regression was used accounting for the binary nature of the response variable (Table 2, Equation 1). For response time, a linear mixed-effects model was fit to the log-transformed data (Table 2, Equation 2). As fixed effects, we entered age, condition, and difficulty into the models. As random effects, we had intercepts for participants and categories. P-values were obtained by likelihood ratio tests of the full model with the effect in question against the model without the effect in question. Post-hoc comparisons were applied using the package emmeans (Lenth et al., 2020).

**Table 2.**
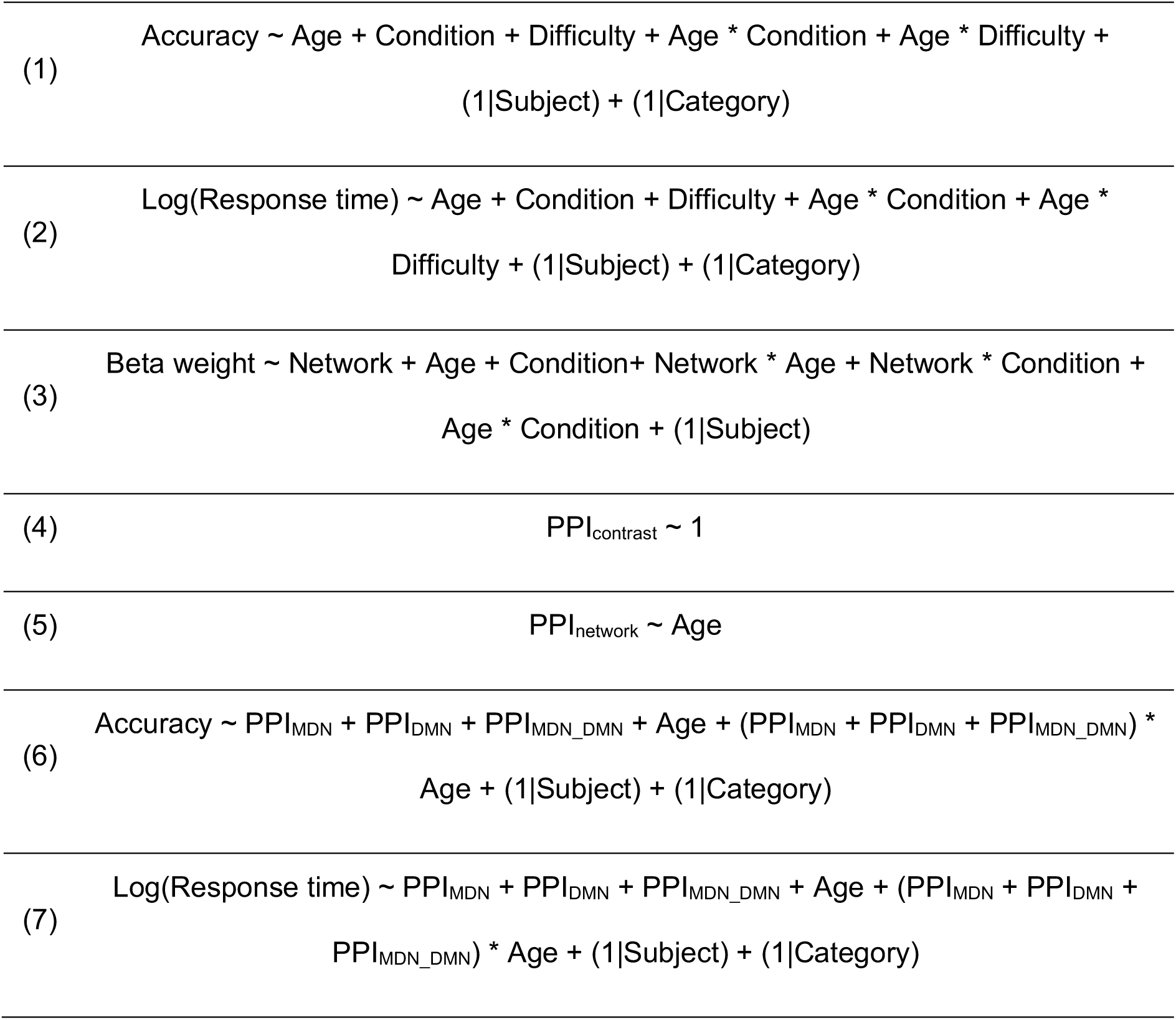
Equations of regression models

#### Functional MRI data

Functional MRI data were modelled in SPM using the two-level approach. On the first level, a general linear model (GLM) was implemented for each participant. The GLM included regressors for the task blocks of the 4 experimental conditions (easy categories, difficult categories, counting forward, and counting backward) and nuisance regressors consisting of the six motion parameters and individual regressors for strong volume-to-volume movement as indicated by values of framewise displacement > 0.9 (Siegel et al., 2014). A two-sample *t*-test indicated that there was no significant difference between older adults (*M* = 15.67, *SD* = 20.04) and young adults (*M* = 7.5, *SD* = 8.35) with respect to the number of regressed volumes (*t*(28.76) = 1.66, *p* = 0.11). Additionally, an individual regressor of no interest was included in the design matrix if a participant had missed a whole task block during the experiment (*n* = 10). Before model estimation, a high-pass filter with a cut-off at 128 s was applied to the data. Statistical parametric maps were generated by estimating the contrast for each condition against rest as well as the direct contrasts between conditions. At the second level, contrast images were entered into a random effects model. For each participant, an averaged mean-centered value of response time was entered as covariate of no interest in the design matrix. For within-group comparisons, one-sample *t*-tests were calculated for the main task-related contrasts Semantic fluency > Counting and Counting > Semantic fluency. To evaluate the modulation of task difficulty within the semantic fluency task, the contrasts Easy > Difficult categories and Difficult > Easy categories were computed.

To investigate the effect of age on task-related activity, we conducted between-group comparisons for the interaction contrasts *c_Age x Semantic fluency_* and *c_Age x Condition_*. Two-sample *t*-tests were carried out using the individual contrast images from the first-level analysis. To ensure that potential areas were indeed active in the respective group, all interactions were characterized by inclusively masking each contrast with significant voxels of the minuend (at *p* < 0.001, uncorr., cf. Noppeney et al., 2006, Meinzer et al., 2012b). A gray matter mask which restricted statistical tests to voxels with a gray matter probability > 0.3 (SPM12 tissue probability map) was applied to all second-level analyses. All results except for the interaction contrasts were corrected for multiple comparisons applying a peak level threshold at *p* < 0.05 with the family-wise error (FWE) method and a cluster-extent threshold of 20 voxels. Interaction results were thresholded at *p* < 0.05 at the cluster level with FWE correction and a voxel-wise threshold at *p* < 0.001. Anatomical locations were identified with the SPM Anatomy Toolbox (Eickhoff et al., 2005, version 2.2c) and the Harvard-Oxford cortical structural atlases distributed with FSL (https://fsl.fmrib.ox.ac.uk). Brain results were rendered by means of BrainNet Viewer (Xia et al., 2013, version 1.7) and MRIcroGL (https://www.mccauslandcenter.sc.edu/mricrogl/, version 1.2.20200331).

Since the strongest activation peaks for both tasks were found in the domain-general systems MDN and DMN, we decided to examine the amount of activation for each condition and age group. We applied binary masks of both networks to analyses within the group of young adults to identify clusters that fell within the respective network. By basing our analysis on activity in the young adults, we ensured that more complex analyses were based on the same number of regions in both age groups. Further, this allowed us to investigate age-related differences in these regions knowing that they are relevant for task processing in young adults. The MDN mask was based on the anatomical parcels of the MD system defined by Fedorenko et al. (2013; available at https://evlab.mit.edu/funcloc/). For the DMN, a mask was created from the 7 networks parcellation by Schaefer et al. (2018). For the contrast Semantic fluency > Counting, peak global and local maxima were found in the MDN, whereas the reverse contrast identified clusters that are typically associated with the DMN. Due to the small number (*n* = 3) of peak clusters for the contrast Counting > Semantic fluency with FWE correction at peak-level, we decided to apply a more lenient threshold (FWE-corrected at cluster-level, *p* < 0.001 at peak-level) for the identification of regions associated with the DMN. This allowed us to extract a similar number of peak maxima for the MDN and the DMN, and provided a much more representative picture of the DMN as a whole. In total, we identified 14 peak maxima in the MDN, and 17 peak maxima in the DMN, respectively (Table 3). Regions of interest (ROIs) for these maxima were created using the MarsBar toolbox (Brett et al., 2002; version 0.44). To this end, identified clusters were extracted from contrast images, spheres of 5 mm from each maxima coordinate were created, and, in a last step, both images were combined. Subsequently, we extracted parameter estimates for these ROIs from the individual contrast images for Semantic fluency > Rest and Counting > Rest. The data were then entered into a linear mixed-effects model with network, age, and condition as fixed effects. A random intercept was included for participants (Table 2, Equation 3). Categorical predictors were sum coded. Significance values were obtained through likelihood ratio tests using the package lme4 (Bates et al., 2015). Post-hoc comparisons were applied using the package emmeans (Lenth et al., 2020).

**Table 3.**
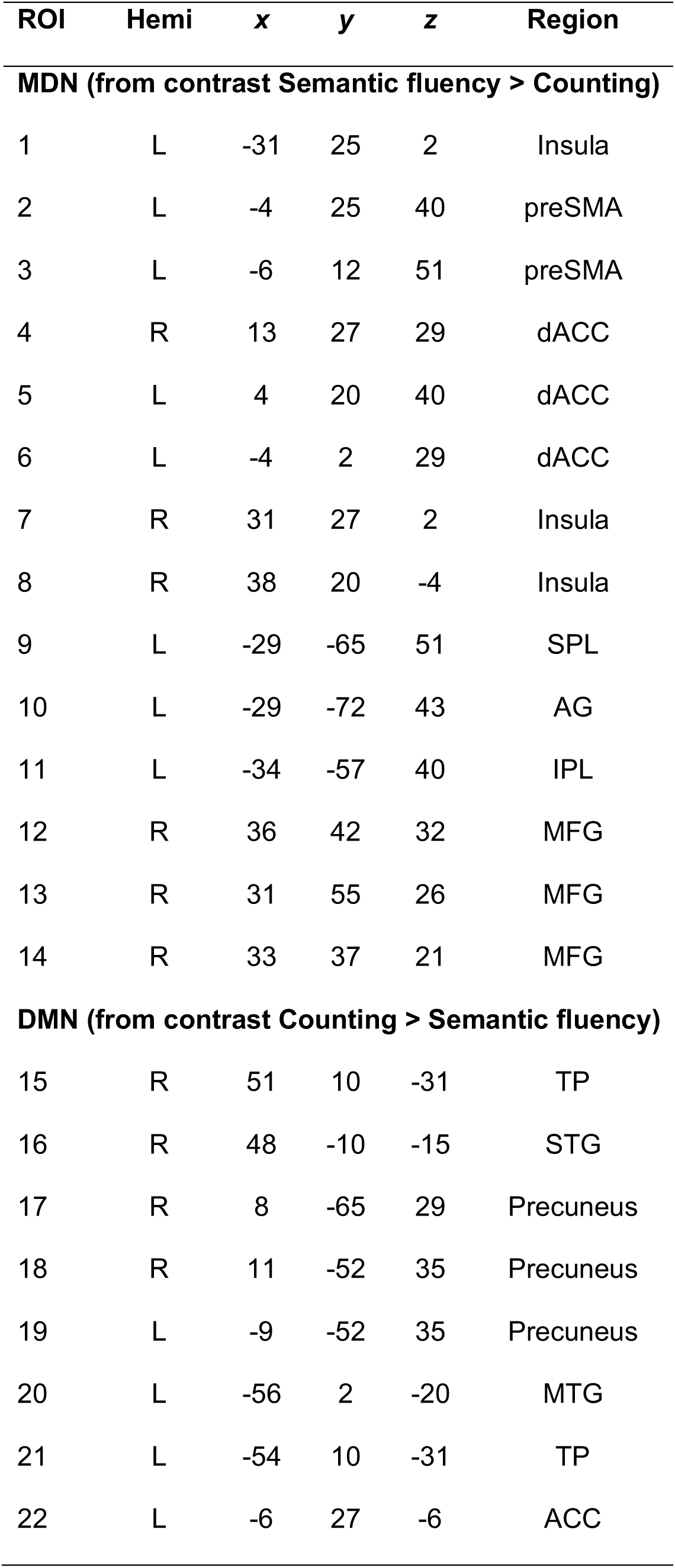
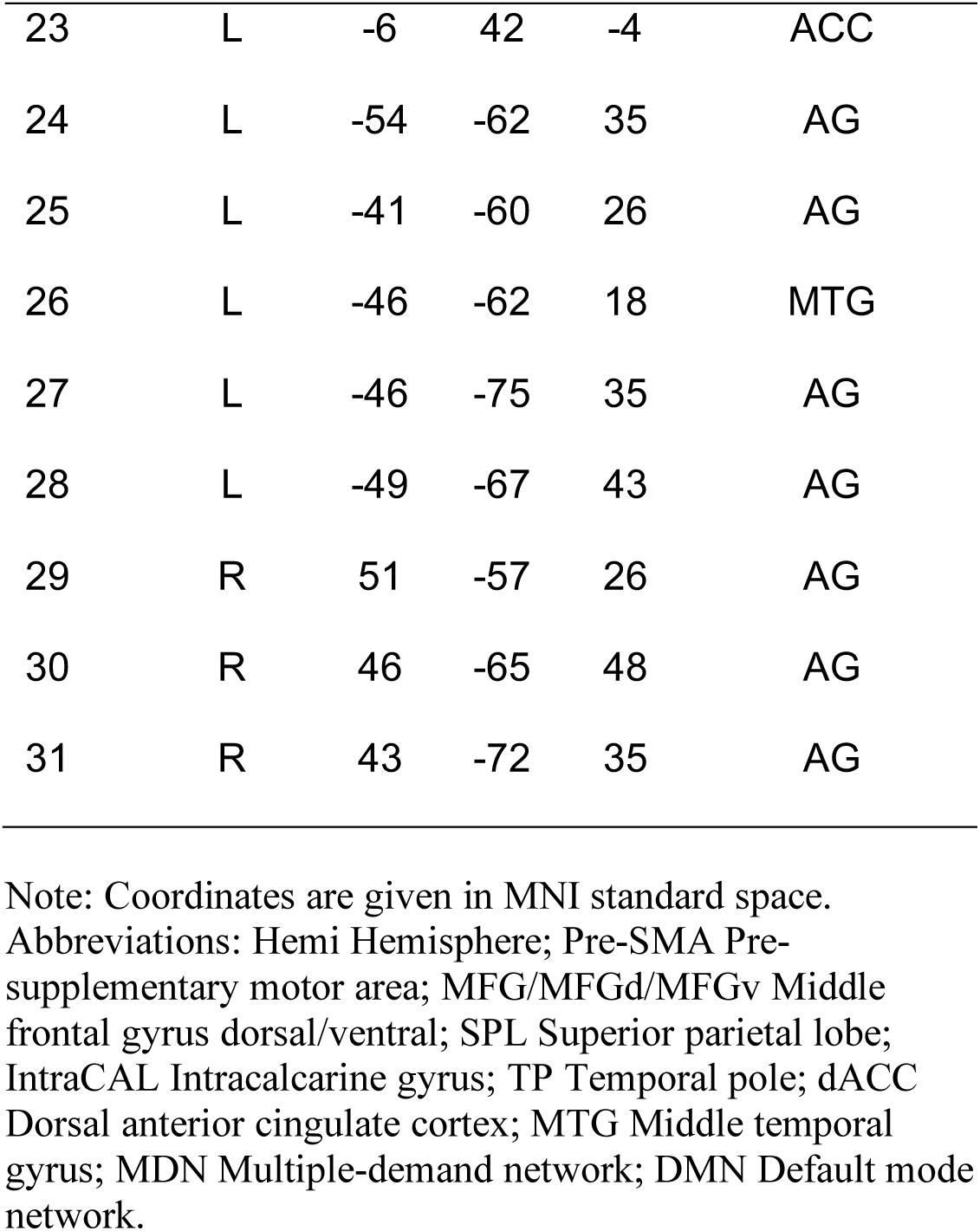
Regions of interest within domain-general networks.

| ROI | Hemi | x | y | z | Region |
| --- | --- | --- | --- | --- | --- |
| **MDN (from contrast Semantic fluency > Counting)** |  |  |  |  |  |
| 1 | L | -31 | 25 | 2 | Insula |
| 2 | L | -4 | 25 | 40 | preSMA |
| 3 | L | -6 | 12 | 51 | preSMA |
| 4 | R | 13 | 27 | 29 | dACC |
| 5 | L | 4 | 20 | 40 | dACC |
| 6 | L | -4 | 2 | 29 | dACC |
| 7 | R | 31 | 27 | 2 | Insula |
| 8 | R | 38 | 20 | -4 | Insula |
| 9 | L | -29 | -65 | 51 | SPL |
| 10 | L | -29 | -72 | 43 | AG |
| 11 | L | -34 | -57 | 40 | IPL |
| 12 | R | 36 | 42 | 32 | MFG |
| 13 | R | 31 | 55 | 26 | MFG |
| 14 | R | 33 | 37 | 21 | MFG |
| **DMN (from contrast Counting > Semantic fluency)** |  |  |  |  |  |
| 15 | R | 51 | 10 | -31 | TP |
| 16 | R | 48 | -10 | -15 | STG |
| 17 | R | 8 | -65 | 29 | Precuneus |
| 18 | R | 11 | -52 | 35 | Precuneus |
| 19 | L | -9 | -52 | 35 | Precuneus |
| 20 | L | -56 | 2 | -20 | MTG |
| 21 | L | -54 | 10 | -31 | TP |
| 22 | L | -6 | 27 | -6 | ACC |
| 23 | L | -6 | 42 | -4 | ACC |
| 24 | L | -54 | -62 | 35 | AG |
| 25 | L | -41 | -60 | 26 | AG |
| 26 | L | -46 | -62 | 18 | MTG |
| 27 | L | -46 | -75 | 35 | AG |
| 28 | L | -49 | -67 | 43 | AG |
| 29 | R | 51 | -57 | 26 | AG |
| 30 | R | 46 | -65 | 48 | AG |
| 31 | R | 43 | -72 | 35 | AG |

Note: Coordinates are given in MNI standard space. Abbreviations: Hemi Hemisphere; Pre-SMA Pre-supplementary motor area; MFG/MFGd/MFGv Middle frontal gyrus dorsal/ventral; SPL Superior parietal lobe; IntraCAL Intracalcarine gyrus; TP Temporal pole; dACC Dorsal anterior cingulate cortex; MTG Middle temporal gyrus; MDN Multiple-demand network; DMN Default mode network.

#### Connectivity analyses

We conducted psychophysiological interaction (PPI) analyses using the gPPI toolbox for SPM12 (McLaren et al., 2012) to investigate task-related modulation of functional connectivity associated with the semantic fluency and the counting task. Furthermore, we applied a modified version of gPPI methods to examine functional connections between individual ROIs during the semantic fluency task (Pongpipat et al., 2020). Seed regions were defined for all previously identified global maxima that were located within the MDN and the DMN (Table 3). For each participant, ROIs were created by searching for the individual peaks within a bounding region of 10 mm relative to the group peak and then drawing a sphere mask (5 mm in diameter) around the individual peak of a given contrast at a threshold of *p* < 0.01. Regression models were set up for each ROI in each participant containing the deconvolved time series of the first eigenvariate of the BOLD signal from the respective ROI as the physiological variable, the four task conditions convolved with the HRF as the psychological variable, and the interaction of both variables as the PPI term. Subsequently, first-level GLMs were calculated. For the gPPI proper methods, contrast images were then entered into a random effects model for group analyses in SPM. We restricted this analysis to the strongest peaks of both contrasts (Semantic fluency > Counting and Counting > Semantic fluency) that fell within the MDN and DMN, respectively. This included the left pre-SMA, bilateral insulae, the right temporal pole and the right precuneus. Our main contrast of interest Semantic fluency > Counting was examined in within-group as well as between-group comparisons by conducting one-sample *t*-tests and a two-sample *t*-test, respectively. Multiple comparison correction was performed with the FWE method at *p* < 0.05 at peak level and a cluster-extent value of 20 voxels. A gray matter mask was applied to all group analyses as described for the task-based fMRI data analysis.

For the modified gPPI, we used the individual first-level GLMs to retrieve parameter estimates (mean regression coefficients). Estimates were extracted for the PPI variable Semantic fluency > Counting for each seed-to-target ROI (1 regression coefficient [PPI] * 31 seed ROIs * 30 target ROIs = 930 parameter estimates per participant). Subsequent group analyses were performed in RStudio (R Core Team, 2018) with the package lme4 (Bates et al., 2015), and visualized using the ggplot2 (Wickham, 2016) and the ggeffects (Lüdecke, 2018) packages. Categorical predictors were sum coded. We were interested in the functional connectivity within and between MDN and DMN regions for each age group. To this end, we calculated intercept-only GLMs where each parameter estimate of each seed and target combination was entered into the model except when seed and target were identical (Table 2, equation 4). The *α*-level (Type I error) for post-hoc comparisons was adjusted using the *Meff* correction (Derringer, 2018). This method estimates the effective number of tests (*Meff*) from the correlations among tested variables and thereby allows for adjusting statistical significance thresholds for multiple comparisons without assuming independence of all tests (Derringer, 2018). The *Meff* value for MDN and MDN variables was calculated to be 27.5. By dividing this value by the overall *α* of 0.05, we obtained a Meff-corrected *α* of 0.0018. For subsequent analyses, the individual parameter estimates of each seed-to-target combination were averaged to create one value per participant for within-MDN, within-DMN, and between-network functional connectivity (3 parameter estimates per participant). To test for an effect of age group on functional connectivity, the parameter estimates were then entered into a GLM with age group as independent variable (Table 2, equation 5). We used Meff-correction to adjust for multiple comparisons. A *Meff* value of 2.49 yielded a Meff-corrected *α* of 0.02.

Furthermore, we were interested in the effect of within- and between-network functional connectivity on participants’ behavioral performance during the in-scanner semantic fluency task. To this end, we calculated generalized mixed-effects logistic regressions for the accuracy data (Table 2, equation 6) and linear mixed-effects models for the log-transformed response time data (Table 2, equation 7). The mean-centered PPI_network_ variables and age group as well as their interaction terms were entered as fixed effects. Random intercepts were included for participants and semantic categories.

Finally, to assess how the observed changes in network properties were related to cognitive performance and semantic memory in general, we performed correlation analyses with the neuropsychological measures that had been tested outside of the scanner. Due to the collinearity of some neuropsychological tests, we first performed a factor analysis on the standardized test scores using maximum likelihood estimation and varimax rotation in RStudio with the package stats (R Core Team, 2018). Based on the hypothesis test (χ^2^ = 14.04, *p* = 0.081) two factors with an eigenvalue > 1 were chosen. For subsequent correlations with functional connectivity measures participant factor scores extracted via regression methods were used.

## Results

### Behavioral Results

For response accuracy, we fitted a generalized linear mixed-effects model for a binomial distribution. Likelihood ratio tests indicated significant main effects of condition (χ^2^ = 21.55, *p* < 0.001) and task difficulty (χ^2^ = 27.43, *p* < 0.001) but not of age group (χ^2^ = 0.19, *p* = 0.66). Further, we detected a significant two-way interaction between age and difficulty (χ^2^ = 9.76, *p* = 0.002) and condition and difficulty (χ^2^ = 3.89, *p* = 0.049) as well as a significant three-way interaction between age, condition, and difficulty (χ^2^ = 9.28, *p* = 0.002). Post-hoc tests applying Bonferroni-corrected pairwise comparisons showed that both age groups produced more correct items in the counting than in the semantic fluency task (all *p* < 0.001) and more items for the easy than the difficult semantic categories (all *p* < 0.001; Fig. 2A; Supplementary Tables S1 and S2).

**Figure 2.**
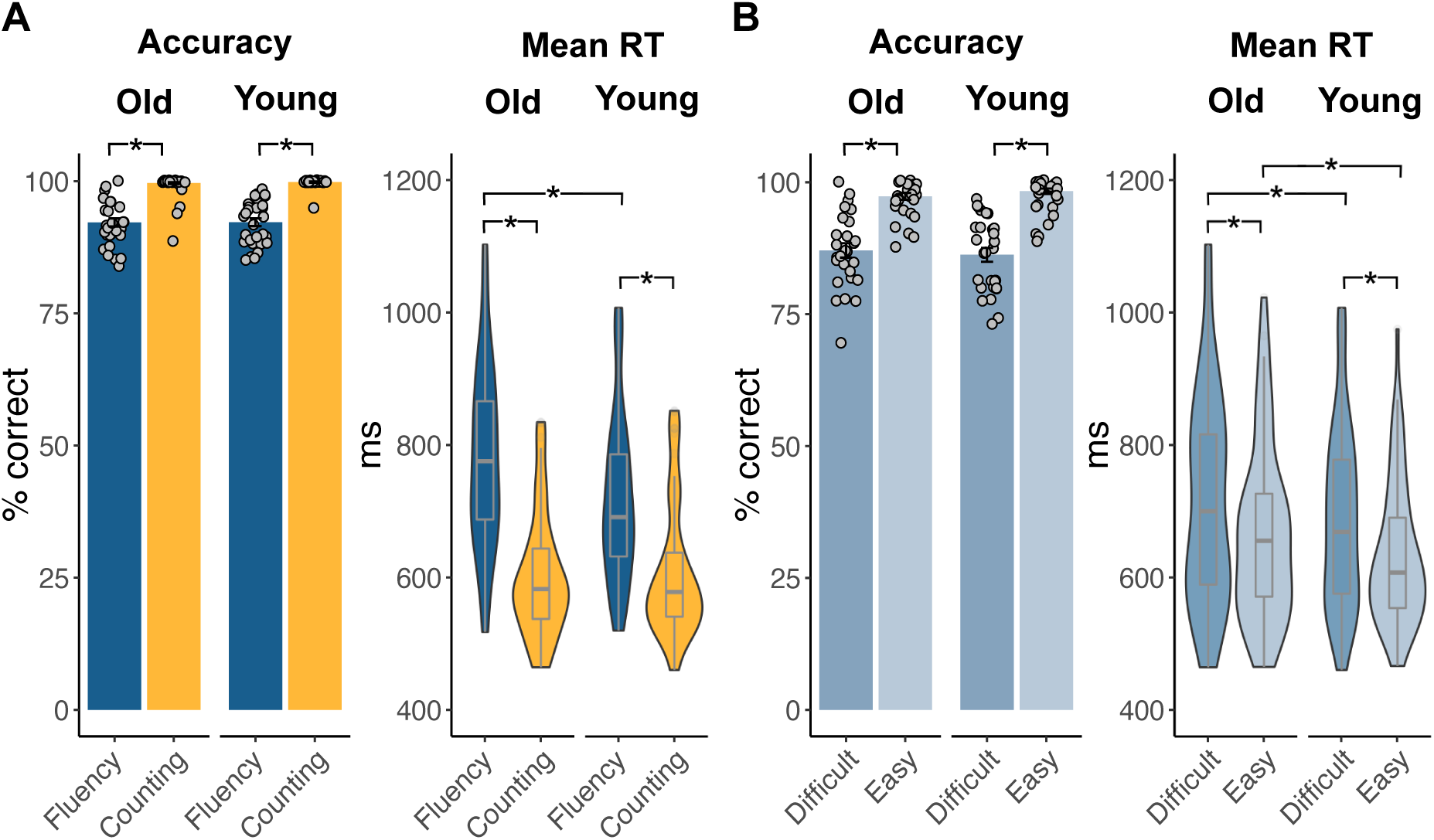
Behavioral results for both age groups. Bar graphs overlaid with mean individual data points for accuracy and violin plots with box plots for mean response times for (A) both tasks (semantic fluency and counting) and (B) difficulty levels (easy and difficult) within semantic fluency. Old: Older adults, Young: young adults. * *p* < 0.001 (Bonferroni-corrected for pairwise comparisons).

Response times were analyzed fitting a linear mixed-effects model after log-transformation of the data. Likelihood ratio tests revealed main effects of age group (χ^2^ = 17.23, *p* < 0.001), condition (χ^2^ = 21.40, *p* < 0.001), and difficulty (χ^2^ = 21.02, *p* < 0.001). There was a significant interaction between age and condition (χ^2^ = 69.46, *p* < 0.001). Post-hoc tests using Bonferroni-corrected pairwise comparisons showed that both age groups responded significantly slower during the semantic fluency task than the counting task and during the difficult than the easy condition (all *p* < 0.001). Furthermore, young adults responded generally faster than older adults during the semantic fluency task (*p* < 0.001), independent of the level of difficulty, but not during the counting task (*p* = 0.05; Fig. 2B; Supplementary Tables S1 and S3).

### Functional MRI Data

#### The effect of task within groups

Both age groups showed similar activation patterns for the main effects of our tasks compared to rest. For semantic fluency, we found a left-lateralized fronto-temporo-parietal network with additional clusters in right frontal and temporal areas, bilateral caudate nuclei, and the cerebellum (Supplementary Fig. S2; Supplementary Tables S4 and S5). The main effect of the less demanding task counting was evident in bilateral activation of sensorimotor cortices and the cerebellum (Supplementary Fig. S2; Supplementary Tables S6 and S7). Within each age group, we were interested in the difference in brain activation between the more demanding semantic fluency task and the automatic speech counting task as well as in the impact of the modulation of task difficulty in the semantic fluency task. For the older adults, the contrast Semantic fluency > Counting revealed a bilateral frontal network with its strongest activation peaks in middle frontal gyri, bilateral insulae extending into inferior frontal gyri, and midline structures comprising superior and medial frontal gyri. Activation in the left hemisphere was further observed in the angular gyrus and superior parietal lobe. Additional bilateral activation peaks were found in the cerebellum, caudate nuclei, calcarine gyri, and thalami (Fig. 3A; Supplementary Table S8). Younger adults demonstrated a similar pattern of activation for the contrast Semantic fluency > Counting, albeit with generally larger clusters in the frontal network (Fig. 3A; Supplementary Table S9). Analyses further yielded separate clusters in the dorsal anterior cingulate cortex (dACC) and the left superior temporal gyrus for the younger group which were not present in the older participants. The reverse contrast (Counting > Semantic fluency) revealed stronger activation in the right hemisphere for both groups. Results showed clusters in the right temporal pole and bilateral precunei (Fig. 3B; Supplementary Tables S10 and S11). In the younger group, additional clusters were observed in bilateral insulae and the middle temporal gyrus. When we applied a more lenient threshold of *p* < 0.001 at peak-level and FWE-correction (*p* < 0.05) at cluster-level, additional peaks were observed in bilateral parietal lobes including angular gyri and the anterior cingulate cortex (Supplementary Table S12).

**Figure 3.**
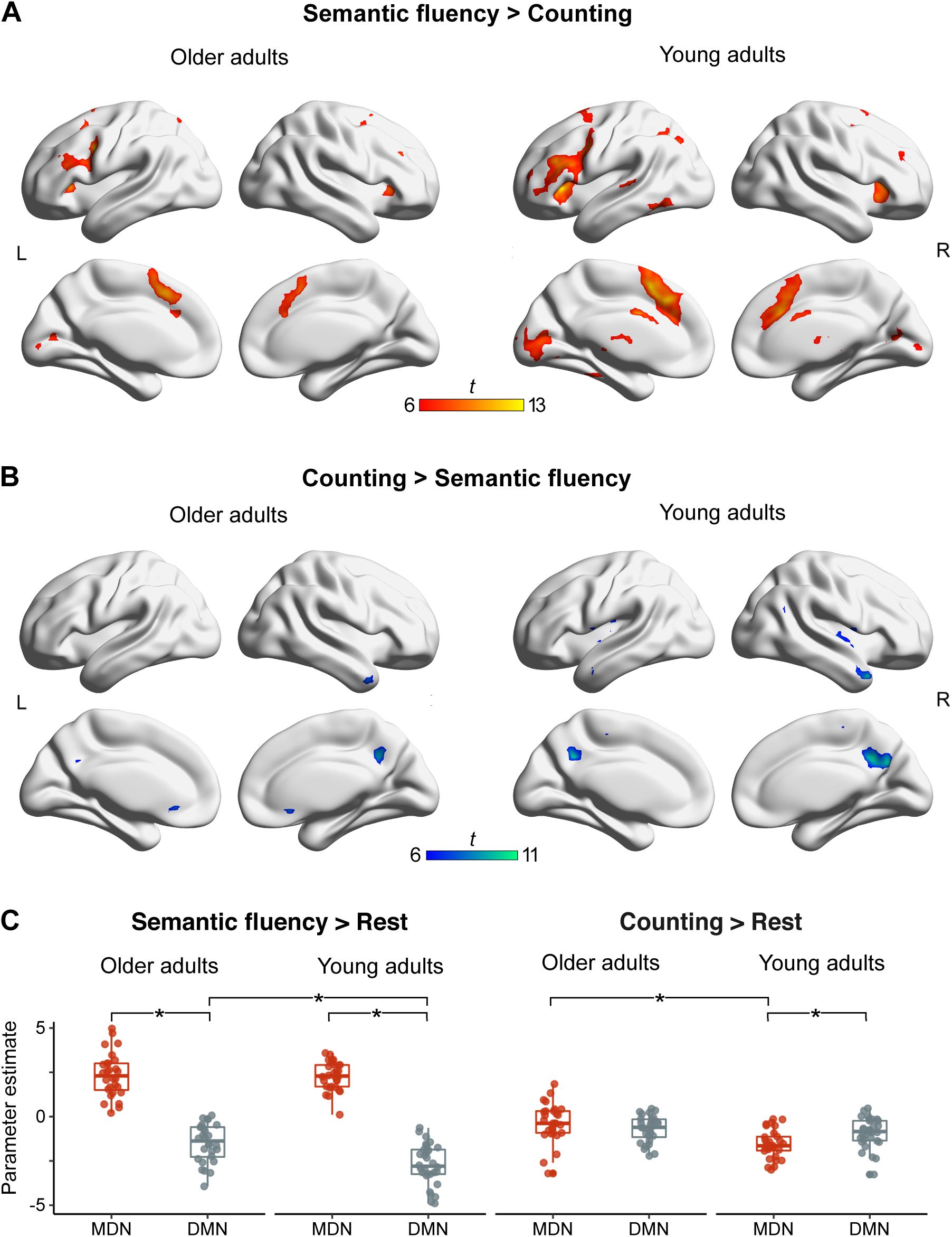
Functional MRI results from univariate analyses for each age group and parameter estimates for peak maxima identified within the MDN and the DMN. (A & B) Results are FWE-corrected at *p* < 0.05 at peak level with a minimum cluster size = 20 voxel. Unthresholded statistical maps are available at https://neurovault.org/collections/9072/. (C) * Significant effects are Bonferroni-corrected. Abbreviations: MDN Multiple-demand network; DMN Default mode network.

A linear mixed-effects model was fit for the mean value of parameter estimates for all peak clusters in the MDN and DMN (Table 3), respectively. Likelihood ratio tests indicated significant effects for the explanatory variables network (χ^2^ = 92.73, *p* < 0.001), age (χ^2^ = 11.03, *p* = 0.017), and condition (χ^2^ = 23.04, *p* < 0.001). We further found a significant interaction between network and condition (χ^2^ = 196.14, *p* < 0.001) and a significant three-way interaction between network, age, and condition (χ^2^ = 18.58, *p* < 0.001). Post-hoc comparisons applying Bonferroni correction revealed an effect of age for the DMN for the Contrast Semantic fluency > Rest with older adults showing stronger activity in DM regions than young adults (*t* = 4.89, *p* < 0.001) as well as an effect of age for the MDN for the Contrast Counting > Rest with older adults showing stronger activity in MD regions than young adults (*t* = 4.81, *p* < 0.001). Moreover, post-hoc tests showed that, in general, the MDN was activated for the semantic fluency task across age groups whereas the DMN showed deactivation (*t* = 24.99, *p* < 0.001). For the counting task, there was no difference in activation between both networks (*t* = 0.81, *p* = 0.42; Fig. 3C; Supplementary Tables S13 and S14).

#### The effect of task difficulty within groups

To investigate the effect of task difficulty on functional brain activation, we contrasted easy and difficult categories from the semantic fluency task in both age groups. We found a significant result only for young adults for the contrast Easy > Difficult categories in the right middle frontal gyrus (Supplementary Table S15).

#### Between-group comparisons

We were interested in the effect of age on task-related activations. For the interaction of both tasks compared to baseline, we found a group difference only during the semantic fluency task for older adults. We detected stronger activity in right frontal regions including superior frontal gyrus and inferior frontal gyrus as well as bilateral parietal lobes (Fig. 4A; Table 4). We were further interested in the interaction of age and condition. The contrast Semantic fluency > Counting revealed a significant interaction with age only for young adults. Stronger activity was observed in the paracingulate gyrus, pre-SMA, and the dACC (Fig. 4B; Table 4). The interaction of age with task difficulty (easy and difficult semantic categories) did not yield any significant results.

**Figure 4.**
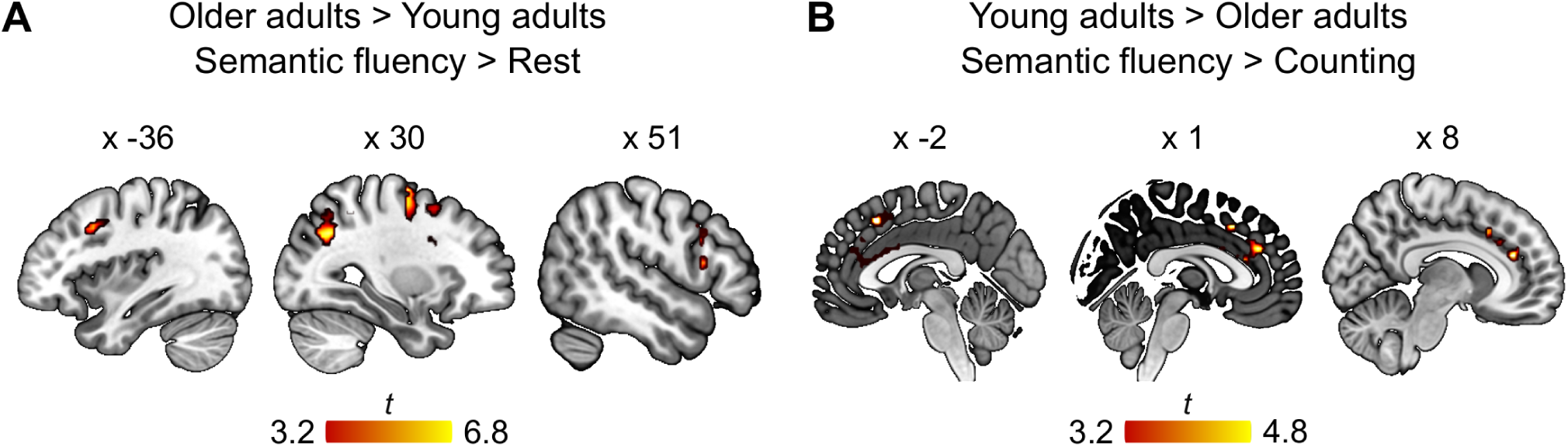
Functional MRI results for interaction effects. Cluster corrected at FWE *p* < 0.05 with a voxel-wise threshold at *p* < 0.001. (A) Restricted to voxels that showed a significant effect of semantic fluency in older adults and (B) restricted to voxels that showed a significant effect of semantic fluency in young adults. Statistical maps are available at https://neurovault.org/collections/9072/.

**Table 4.** Results for age-dependent differences in task-related activity.

| Anatomical structure | Hemi | *k* | *t* | *x* | *y* | *z* |
| --- | --- | --- | --- | --- | --- | --- |
| Interaction older > young adults for semantic fluency > rest (inclusively masked with [older adults: semantic fluency > rest]) |  |  |  |  |  |  |
| **Superior frontal gyrus** | **R** | **110** | **6.84** | **28** | **-8** | **65** |
| Precentral gyrus | R |  | 5.53 | 28 | -8 | 54 |
| **Superior parietal lobe** | **R** | **263** | **6.82** | **18** | **-60** | **60** |
| Middle occipital gyrus | R |  | 6.04 | 31 | -62 | 35 |
| Precuneus | R |  | 5.93 | 11 | -55 | 60 |
| Precuneus | R |  | 5.39 | 13 | -67 | 62 |
| **Middle frontal gyrus** | **L** | **74** | **5.38** | **-36** | **15** | **38** |
| Middle frontal gyrus | L |  | 4.05 | -46 | 17 | 38 |
| Precentral gyrus | L |  | 3.76 | -41 | 2 | 48 |
| **Inferior parietal lobe** | **L** | **109** | **5.12** | **-31** | **-45** | **54** |
| Superior parietal lobe | L |  | 4.32 | -19 | -60 | 46 |
| Precuneus | L |  | 4.28 | -11 | -70 | 48 |
| Superior parietal lobe | L |  | 4.19 | -16 | -62 | 60 |
| **Middle frontal gyrus** | **R** | **132** | **4.96** | **36** | **5** | **38** |
| Inferior frontal gyrus, p.tr. | R |  | 4.53 | 36 | 17 | 24 |
| Inferior frontal gyrus, p.op. | R |  | 4.07 | 48 | 17 | 13 |
| Inferior frontal gyrus, p.op. | R |  | 3.73 | 51 | 15 | 26 |
| **Middle frontal gyrus** | **R** | **74** | **4.47** | **26** | **12** | **51** |
| **Interaction young > older adults for semantic fluency > counting** (inclusively masked with [young adults: semantic fluency > counting]) |  |  |  |  |  |  |
| **Pre-SMA** | **L** | **79** | **4.84** | **-1** | **20** | **46** |
| Pre-SMA | R |  | 3.81 | 11 | 27 | 32 |
| Pre-SMA | L |  | 3.55 | -1 | 10 | 48 |
| **Anterior cingulate cortex** | **R** | **114** | **4.58** | **8** | **37** | **21** |
| Pre-SMA | R |  | 4.36 | 1 | 37 | 26 |
| Anterior cingulate cortex | L |  | 4.13 | -1 | 20 | 21 |
| Anterior cingulate cortex | L |  | 3.58 | -4 | 5 | 29 |

Note: Cluster corrected at FWE $p < 0.05$ with a voxel-wise threshold at $p < 0.001$ . Coordinates are given in MNI standard space, cluster size (k) is given in $\text{mm}^3$ . Note that no significant differences above cluster correction threshold were found for (i) interaction young > older adults for semantic fluency > rest, (ii) interaction older > young adults for semantic fluency > counting. Abbreviations: Hemi Hemisphere; p. tr. pars triangularis; p.op. pars opercularis; pre-SMA pre-supplementary motor area.

### Generalized psychophysiological interactions

Based on the activation patterns from our univariate within-group analyses, we conducted traditional gPPI analyses for the 5 strongest activation peaks that fell within the MDN or DMN. We asked whether and how increased semantic task demands modulate connectivity of our ROIs.

#### Whole-brain functional connectivity for semantic fluency

Three ROIs were extracted from the univariate contrast Semantic fluency > Counting, the left pre-SMA and bilateral insulae. For the seeds in the left pre-SMA and left insula, analyses revealed only significant clusters in the group of younger adults, whereas the seed in the right insula yielded only significant results for the older adults. The left pre-SMA showed increased connectivity with subcortical structures (bilateral caudate nuclei and thalami) as well as with the left precuneus and PCC in the parietal lobe (Fig. 5A; Supplementary Table S16). For the seed in the left insula, we found a similar connectivity pattern. Significant coupling was observed with bilateral caudate nuclei and the left precuneus (Fig. 5B; Supplementary Table S17). For the older adults, the right insula showed significant coupling with the precuneus and pars orbitalis in left inferior frontal gyrus (Fig. 5C; Supplementary Table S18).

**Figure 5.**
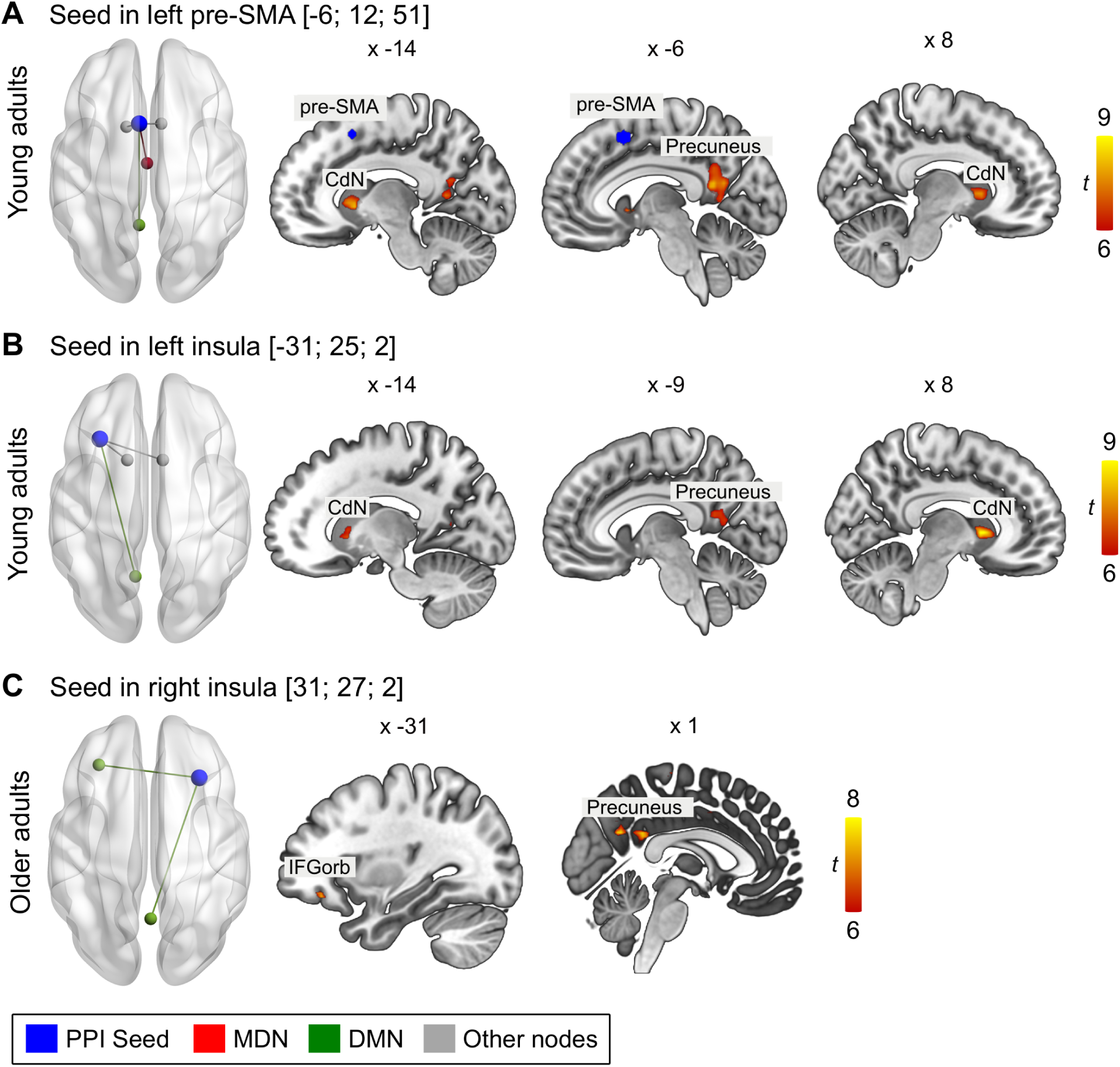
Functional connectivity for seeds from contrast Semantic fluency > Counting. All results are FWE-corrected at *p* < 0.05 at peak level with a minimum cluster size = 20 voxel. Abbreviations: Pre-SMA presupplementary motor area; CdN caudate nucleus; IFGorb inferior frontal gyrus, pars orbitalis. Unthresholded statistical maps are available at https://neurovault.org/collections/9072/.

Moreover, two ROIs from the contrast Counting > Semantic fluency which were associated with the DMN, the right temporal pole and the right precuneus, were used for traditional gPPI analyses. Seeding in the right temporal pole showed increased functional connectivity exclusively in the ipsilateral hemisphere. For the older adults, we found a significant cluster in inferior frontal gyrus (pars opercularis) which extended into the insula (Fig. 6A; Supplementary Table S19). For the young adults, results revealed significant clusters in the inferior frontal gyrus (pars opercularis), superior frontal gyrus, insula, and supramarginal gyrus (Fig. 6A, Supplementary Table S19). The seed in the right precuneus revealed extensive bilateral functional coupling in both age groups. For the older adults, the right precuneus showed prominent connectivity with frontal, temporal, and parietal areas in both hemispheres (Fig. 6B; Supplementary Table S20). A similar pattern emerged for the group of younger adults, albeit with a greater number of significant clusters (Fig. 6B; Supplementary Table S20). Two-sample *t*-tests did not show significant differences between groups in the PPI results for either task.

**Figure 6.**
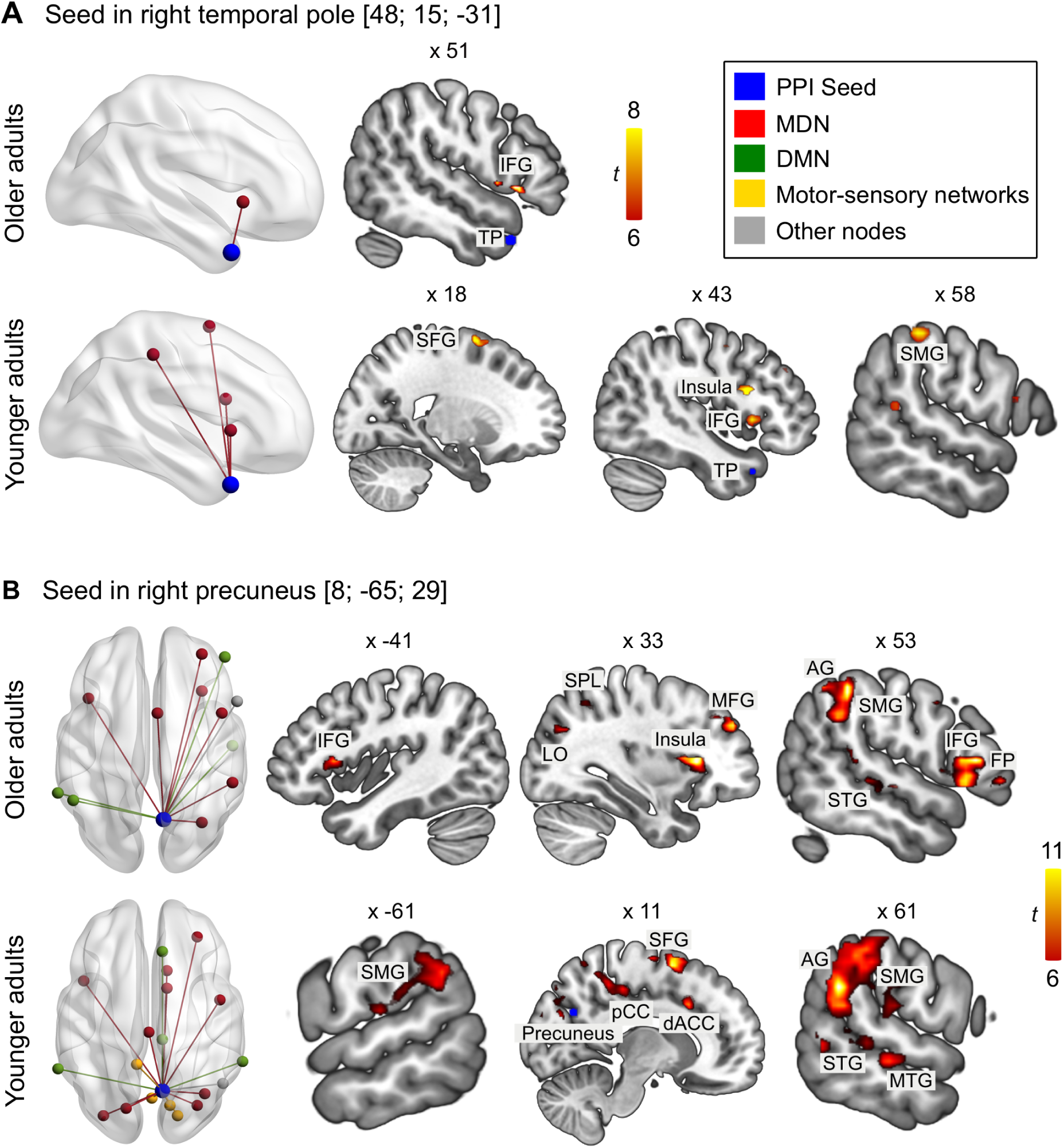
Functional connectivity for seeds from contrast Counting > Semantic fluency. All results are FWE-corrected at *p* < 0.05 at peak level with a minimum cluster size = 20 voxel. Abbreviations: IFG inferior frontal gyrus; TP temporal pole; SFG superior frontal gyrus; SMG supramarginal gyrus; LO lateral occipital cortex; SPL superior parietal lobe; MFG middle frontal gyrus; AG angular gyrus; FP frontal pole; STG superior temporal gyrus; pCC posterior cingulate cortex; dACC dorsal anterior cingulate cortex; MTG middle temporal gyrus. Unthresholded statistical maps are available at https://neurovault.org/collections/9072/.

#### Within- and between-network functional connectivity during semantic fluency

To further examine the task-related connectivity within and between the domain-general systems MDN and DMN during the semantic fluency task compared to counting, we conducted modified gPPI analyses. For each seed-to-target combination of the ROIs in MDN and DMN (Table 3), we calculated intercept-only GLMs for each age group (Fig. 7A). For older adults, the results showed significant positive functional connectivity for regions within the MDN but not for regions within the DMN. Further, the analyses revealed strong coupling for regions between MD and DM networks. A similar pattern was observed for young adults, albeit with overall stronger connectivity. Compared to the counting task, results showed strengthened functional connectivity within regions of the MDN and for regions between MD and DM networks during semantic fluency.

**Figure 7.**
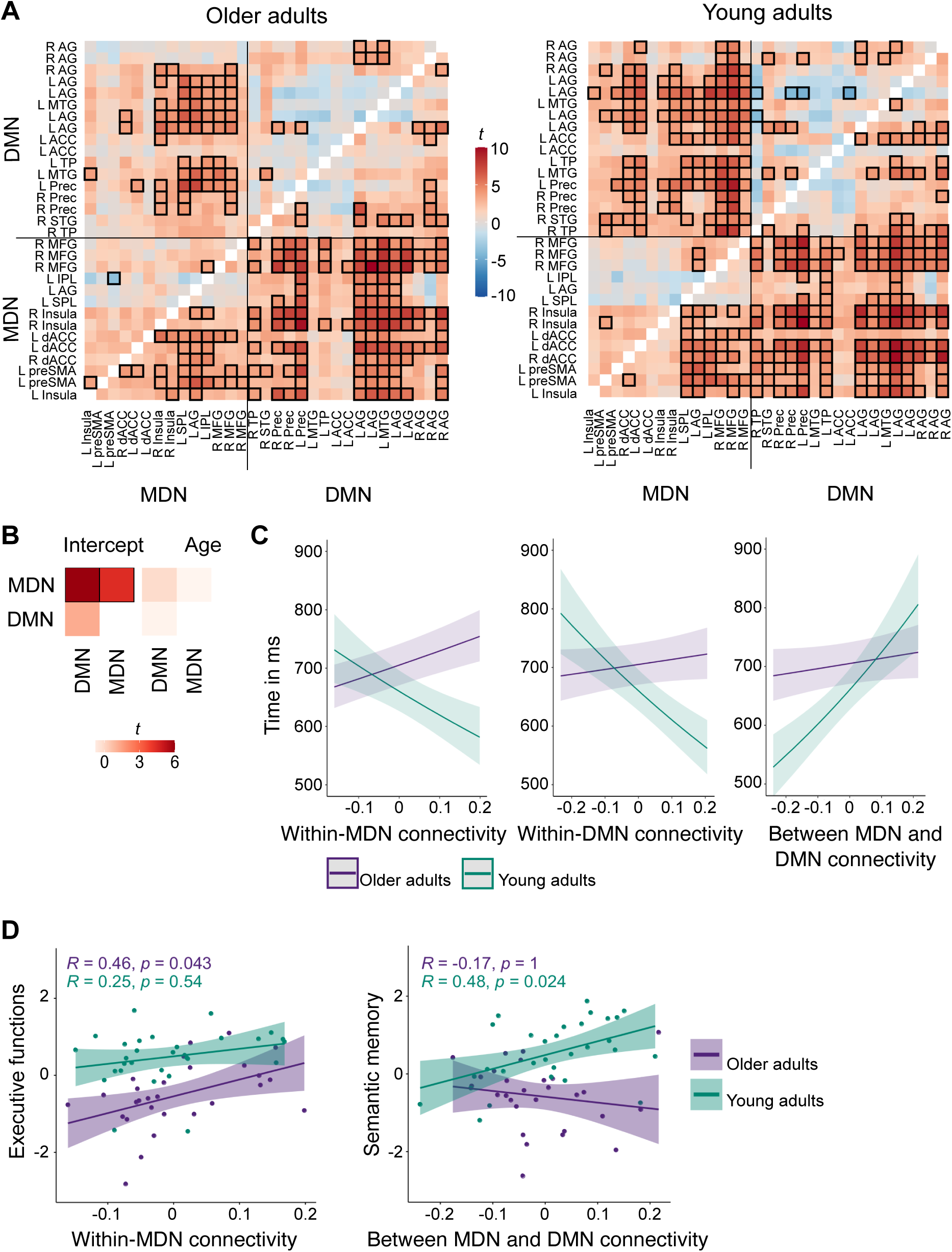
Functional connectivity of domain-general network regions during semantic fluency. (A) Within- and between-network functional connectivity for each seed to target combination for each age group. Heatmaps show *t* values. Thresholded *t* values with Meff-corrected *α* of 0.0018 are indicated with black boxes. (B) Effect of age on functional connectivity. There were no age differences in functional connectivity between or within networks. Heatmaps show *t* values with Meff-corrected *α* of 0.02. Significant effects are indicated with black boxes. (C) Significant two-way interactions between age and functional connectivity for response time during semantic fluency. (D) Correlation analyses between functional connectivity measures and neuropsychological factors. For within-MDN connectivity, only older adults showed a significant correlation with executive functions, while only young adults showed a significant correlation between semantic memory and between-network functional connectivity. Abbreviations: R right hemisphere; L left hemisphere; AG angular gyrus; MTG middle temporal gyrus; ACC anterior cingulate cortex; TP temporal pole; Prec Precuneus; STG superior temporal gyrus; MFG middle frontal gyrus; IPL inferior parietal lobe; SPL superior parietal lobe; dACC dorsal anterior cingulate cortex; preSMA presupplementary motor area; MDN Multiple-demand network; DMN Default mode network.

#### Effect of age on within- and between-network functional connectivity

We were interested whether there was an effect of age group on the within- and between network functional connectivity. To this end, each PPI network pair (within-MDN, within-DMN, and between MDN and DMN) was regressed on age (Fig. 7B; Supplementary Table S21). Multiple-comparisons correction was performed using Meff-correction. The results did not show a significant effect of age on within- and between-network functional connectivity (*p*’s > 0.3) suggesting that the strength of functional connectivity was age-invariant.

#### Effect of functional connectivity on in-scanner task performance

To determine whether functional connectivity within and between regions of MDN and DMN predicted participant’s in-scanner task performance, we fitted generalized mixed-effects models for accuracy and response time as outcome variables and functional connectivity, age, and their interaction terms as explanatory variables. Since the functional connectivity measures were based on our contrast of interest Semantic fluency > Counting, statistical models were only fit for the behavioral results for the semantic fluency and not for the counting task. The results did not indicate significant effects of functional connectivity on accuracy (Supplementary Table S22). However, analyses revealed significant effects of functional network connectivity on response time (Fig. 7C; Supplementary Table S22 and S23). We identified main effects of within-DMN (χ^2^ = 12.65, *p* < 0.001), and between-network functional connectivity (χ^2^ = 29.43, *p* < 0.001) as well as of age (χ^2^ = 63.15, *p* < 0.001). Beta coefficients indicated that across networks strengthened functional connectivity was associated with slower response times and that young adults performed generally faster than older adults which confirmed our behavioral results. Moreover, significant interactions between age and within-MDN (χ^2^ = 32.84, *p* < 0.001), within-DMN (χ^2^ = 37.33, *p* < 0.001), and between-network functional connectivity (χ^2^ = 24.78, *p* < 0.001) were found. Post-hoc tests showed that age group had a different effect on within- and between-network functional connectivity. While response time increased with strengthened connectivity in the MDN and the DMN for older adults, the opposite pattern was observed for young adults who responded faster when within-network functional connectivity increased (all *p* < 0.001). For functional connectivity between MDN and DMN, stronger coupling predicted slower responses in both age groups, albeit with young adults showing a significantly steeper positive slope than older adults (*p* < 0.001; Fig. 7C).

#### Effect of functional connectivity on cognitive performance and semantic memory

We were interested in the relationship between functional connectivity and general cognitive and semantic memory performance which were assessed via neuropsychological tests outside of the scanner. Since some tests showed high collinearity, we first performed a factor analysis on the data of both age groups together. Results identified two factors: A *cognitive performance* factor with high loadings on trail making tests A (0.8) and B (0.71), the digit symbol substitution test (0.73), and the reading span test (0.45), and a *semantic memory* factor with high loadings on the spot-the-word test (0.5) and the two verbal fluency tests for hobbies (0.44) and surnames (0.98). Individual factor scores for participants were extracted and subsequently correlated with functional connectivity measures. The resulting *p*-values were corrected for multiple comparisons using Bonferroni-correction (*p* = 0.05/3 functional connectivity parameters = 0.017). First, we used partial Pearson correlations to test for a relation between connectivity and cognitive performance while controlling for the effect of age. Results revealed a significant positive correlation between executive functions and within-MDN functional connectivity (*r* = 0.36, *p* = 0.018). Second, we calculated Pearson correlations within each age group. For older adults, we found a significant positive correlation between cognitive performance and within-MDN functional connectivity (*r* = 0.46, *p* = 0.043; Fig. 7D). For young adults, results showed a significant positive correlation between semantic memory and functional connectivity between MDN and DMN (*r* = 0.48, *p* = 0.024; Fig. 7D).

Taken together, functional connectivity within- and between-MD and DM network regions was associated with efficiency during the experimental task as well as general cognitive and semantic performance in both age groups. The effect of functional connectivity on response time was moderated by age with young adults profiting from a strengthened within-network connectivity whereas older adults showed a decline in response speed. Furthermore, both age groups performed slower when functional connectivity between both domain-general systems increased. Finally, functional connectivity was differently related to out-of-scanner tasks in both age groups. While analyses revealed a positive association between cognitive performance and within-MDN functional connectivity in older adults, between-network functional connectivity showed a positive effect on semantic memory in young adults.

## Discussion

The current study set out to describe the effects of aging on the interplay of domain-specific and domain-general neural networks in semantic cognition. By contrasting a semantic fluency task with a low-level verbal control task in an fMRI experiment, we delineated two distinct task-related networks which displayed strong overlap with the domain-general MD and DM systems. Using task-based connectivity analyses, our findings point towards a strong interaction of these networks during verbal semantic processing across age groups and lend support to the notion that integration between usually anti-correlated functional networks increases for tasks that require cognitive control (Shine et al., 2016). Importantly, our results provide new insights into the impact of age on the functional coupling within and between MDN and DMN regions when semantic knowledge is retrieved in a goal-directed manner from memory. In line with a recent suggestion that additional recruitment of the prefrontal cortex in older adulthood might not reflect compensation but rather reduced efficiency or specificity (Morcom and Henson, 2018), we show here that increased in-phase synchronization of task-relevant networks is generally associated with a decline in task efficiency in older adults whereas young adults capitalize more on strengthened functional connectivity. This finding sheds new light on the frequently reported pattern of strengthened between-network functional connectivity in older adults at rest (Chan et al., 2014; Geerligs et al., 2015; Spreng et al., 2016).

Our task paradigm revealed two distinct functional networks for semantic fluency and counting. The main effect of semantic fluency displayed a predominantly left lateralized fronto-temporo-parietal network for both age groups with additional activation peaks in right frontal and temporal areas, bilateral caudate nuclei, and the cerebellum. These results align well with previous investigations that applied a semantic fluency paradigm (Baciu et al., 2016; Birn et al., 2010; Marsolais et al., 2014; Meinzer et al., 2009, 2012a, 2012b; Nagels et al., 2012; Vitali et al., 2005; Wagner et al., 2014; Whitney et al., 2009). The main effect of the counting task was evident in both groups in bilateral activation of sensorimotor cortices and the cerebellum which is consistent with previous studies that used an automated speech task (e.g., Birn et al., 2010; Geranmayeh et al., 2014; Marsolais et al., 2014). Further, older adults showed recruitment of the pre-supplementary motor area which could reflect increased cognitive demands for this age group while keeping track of the numbers during counting. In the direct comparison of both tasks, semantic fluency elicited a network that resembled the main effect of the task minus activity in pre- and postcentral gyri in both age groups which corroborates the functional role of this network in spoken language beyond low-level sensorimotor aspects (Geranmayeh et al., 2014). Significant activation for the counting task compared to semantic fluency was evident in a mainly right lateralized network. Previous studies have suggested that neural networks for highly overlearned automated speech tasks are either right-lateralized in healthy participants (Sidtis et al., 2009; Vanlancker-Sidtis et al., 2003) or show less left lateralization than semantically rich language production tasks (Bookheimer et al., 2000; Petrovich Brennan et al., 2007). Further evidence stems from the common observation that automated speech (e.g., counting) is often preserved in patients who suffer from aphasia after a left hemisphere stroke (Vanlancker-Sidtis et al., 2003).

Despite the semantic nature of the task, we found that the strongest activation clusters for semantic fluency were located in the domain-general MD system in both age groups. These results are in line with previous studies that applied a similar task (e.g., Basho et al., 2007; Lurito et al., 2000) and highlight the strong executive aspect of this paradigm. There is emerging evidence on the overlap of language-specific regions like the left inferior frontal gyrus (IFG) with networks that are implicated in domain-general executive processing (Fedorenko et al., 2012) and semantic control processes (Jackson et al., 2021; Noonan et al., 2013; Thompson-Schill et al., 1997). A recent meta-analysis demonstrated an overlap of some regions of the semantic control network with the MDN thus emphasizing the role of domain-general control in language processing (Jackson, 2021). Here, we observed that semantic fluency predominantly activated the domain-general regions of the semantic control network, like the pre-SMA and the dorsomedial prefrontal cortex including dorsal IFG, and only a small part of domain-specific semantic control (ventral IFG). Hence, the domain-general control regions may not be language-specific but appear to strongly contribute to a task that requires goal-directed controlled access to semantic memory while monitoring the verbal articulation of words that match the semantic categories. The scope of the present investigation was confined to the age-dependent contribution of domain-general systems to semantic cognition. Nonetheless, their interaction with the semantic network remains certainly an important question for future research. Further support for the contribution of executive functions to semantic fluency stems from behavioral studies that associated cognitive flexibility, inhibition, working memory, and attention with successful performance (Aita et al., 2018; Amunts et al., 2020; Gordon et al., 2018). For both age groups, peak clusters of counting were found in the posterior DMN which is in line with our expectation of a low-level language production task in comparison to the more demanding semantic fluency task.

Our whole-brain functional connectivity results based on traditional gPPI analyses showed that regions in the domain-general MD and DM systems strongly interact during a semantic word retrieval task compared to counting across both age groups. This was true for seeds coming from the MDN as well as the DMN. This finding is in line with previous studies that reported task-specific functional coupling of cognitive control regions with the DMN, most notably the posterior cingulate cortex (PCC)/precuneus, especially in tasks requiring controlled access to semantic memory (Krieger-Redwood et al., 2016; Smith et al., 2016). Remarkably, the PCC/precuneus was the only region in our study that showed functional connectivity with all seeds from the MDN and displayed extensive functional coupling with multiple nodes in the DMN as well as other neural networks in both age groups. This finding stresses its role as a cortical hub connecting networks to support complex behavior (Leech et al., 2012). The functional coupling of MDN and DMN is especially interesting in light of our functional results where we observed significant deactivation in regions of the DMN during semantic fluency in both age groups. It corroborates the notion that networks that are anti-correlated during task can still show functional integration in contextually relevant situations to facilitate goal-directed behavior (Krieger-Redwood et al., 2016; Spreng et al., 2014).

We gained further insight into the task-related functional integration of MD and DM network regions by our analyses of phase synchronization within and between both domain-general systems. Our results show that functional coupling within the MDN and between the MDN and the DMN strengthened with an increase of task load, which was true independent of age. First, the positive coupling within regions of the MDN is in line with our univariate results for semantic fluency: here, we found that the strongest activation clusters were located in the MDN in both age groups, thus confirming the necessary engagement of this network for successful task performance. Second, the strong in-phase synchronicity between regions of the MD and DM network for semantic fluency compared to the control task complement our PPI results which showed a strong interaction of both domain-general systems. This is line with the notion that the integration of the DMN is relevant for successful task processing in memory-guided cognition (Smith et al., 2016; Vatansever et al., 2015), especially when access to semantic memory is required (Wirth et al., 2011; Krieger-Redwood et al., 2019).

Interestingly, our results on whole-brain as well as within- and between-network functional connectivity did not reveal an effect of age. There is an extant literature describing age-related changes in connectivity in resting state networks with the most common observation of decreased within- and increased between-network functional connectivity (Chan et al., 2014; Ferreira et al., 2016; Geerligs et al., 2015; Grady et al., 2016; Ng et al., 2016; Zonneveld et al., 2019). However, results are more inconsistent for task-related changes in functional connectivity with age. Across a range of cognitive tasks, studies reported a similar pattern as for resting state investigations (Geerligs et al., 2014; Spreng et al., 2016), no changes for within-but only for between-network connectivity (Gallen et al., 2016; Grady et al., 2016), as well as age invariance (Pongpipat et al., 2020; Trelle et al., 2019). In the domain of semantic cognition, findings are sparse with one study observing reduced within-network integration for a semantic fluency task which was not associated with a poorer performance in older adults (Marsolais et al., 2014). There are two possible explanations for the present age invariance in functional connectivity. First, the group of older adults in our study might have been too young to detect changes in functional connectivity. Longitudinal studies on cognitive aging showed that a turning point in functional coupling takes place around the age of 65-70 years (Ng et al., 2016; Zonneveld et al., 2019). Thus, although the young adults in our study displayed numerically greater and stronger functional coupling than the older adults, the overall pattern was too similar in both groups. Second, the lack of an age effect on functional connectivity might be related to the semantic nature of our fluency paradigm. Semantic tasks have been shown to require functional coupling between cognitive control as well as DM regions like the PCC/precuneus which has been implicated in semantic cognition even in young adults (Krieger-Redwood et al., 2016). Hence, this might have aggravated the possibility of observing the frequently reported increase of between-network functional connectivity in older adults and underlines the necessity for more task-based investigations in the future to better understand the picture of neurocognitive aging.

Intriguingly, despite the observed age invariance, functional connectivity had different effects on in-scanner task performance and cognitive functioning in both age groups. Our results show that older adults did not capitalize on strengthened functional connectivity in the same way as young adults. This was the case for functional connectivity within the MDN and the DMN where an increase of connectivity was associated with slower performance in the semantic fluency task in older adults but with faster performance in young adults (Fig. 8A). In contrast, strengthened between-network functional connectivity led to a slower performance in both age groups, with a stronger effect for young adults (Fig. 8B). Considering our whole-brain connectivity results that showed strong positive coupling between both networks during semantic fluency, this decrease in efficiency might reflect the more effortful communication between task-relevant networks compared to within-network coupling, hence leading to a slower performance across age groups. Interestingly, despite the negative effect on task efficiency, increased functional coupling between MD and DM had a positive effect on semantic memory in young but not in older adults. For the latter group, strengthened connectivity within MD regions was associated with better cognitive performance, albeit still at a significantly lower level of performance than in young adults.

**Figure 8.**
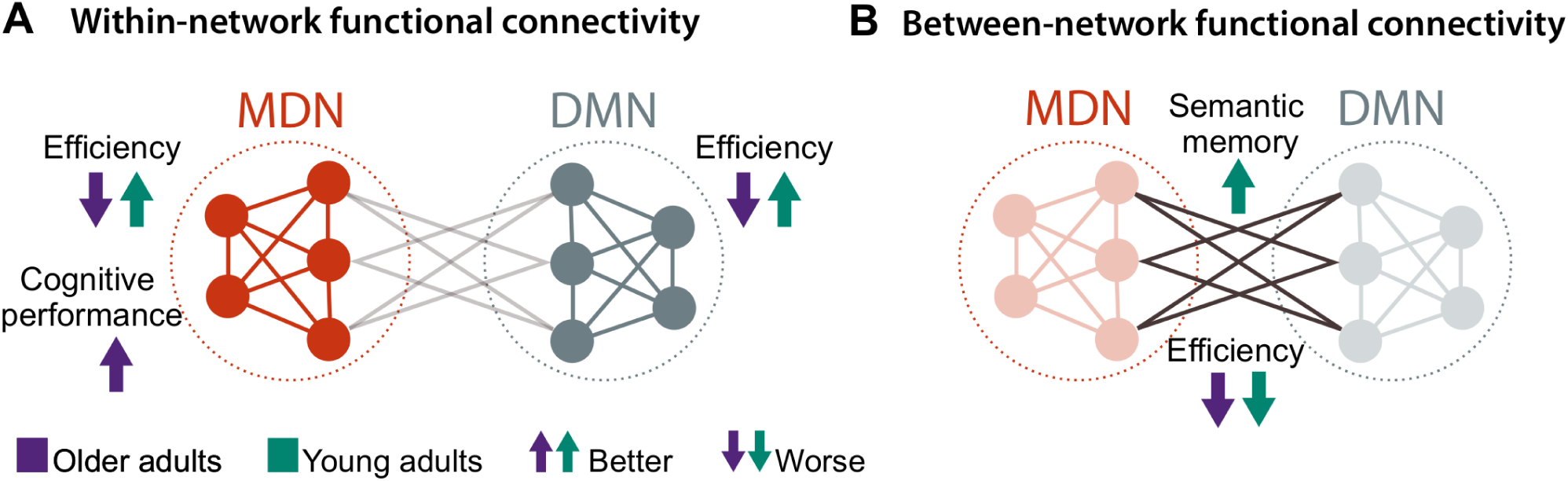
The different effects of within- and between network functional connectivity on task performance in each age group. (A) Young adults improved their efficiency in the form of faster response times whereas older adults performed slower when functional connectivity within the MDN or DMN increased. Moreover, strengthened connectivity within the MDN was related to a better performance in executive measures for older adults. (B) Strengthened between-network functional connectivity led to a decline in efficiency in both age groups. However, it was also associated with an improved performance in semantic memory only for young adults. Abbrev.: MDN Multiple-demand network; DMN Default mode network.

Overall, our findings on the age-dependent relevance of functional connectivity to behavior are in line with theories of neurocognitive aging that suggest a reduced efficiency of neural networks with age (Davis et al., 2012; Geerligs et al., 2014; Shafto and Tyler, 2014). Although older adults rely on similar neural networks as young adults for task processing, they cannot equally capitalize on them. Young adults increased their performance as well as their processing efficiency with strengthened in-phase synchronization of task-relevant networks while older adults showed improvements in cognitive performance but not in efficiency with increased connectivity. Our results thus provide new insights into the behavioral relevance of the frequently observed pattern of neural dedifferentiation (Baltes and Lindenberger, 1997; Grady, 2012; Park et al., 2004) showing that older adults do not engage task-relevant networks in the same beneficial way as young adults. This is especially relevant in the context of semantic cognition where, according to the DECHA framework, increased semantic knowledge with age could lead to a performance advantage (Spreng and Turner, 2019). Here, we show that this is not the case in a task that requires an efficient use of control systems while accessing semantic memory; thus, lending support to the notion that older adults are less flexible in the goal-directed functional coupling of executive and default resources (Spreng and Turner, 2019).

Our findings on age-related differences in cortical activation for both tasks further underline the observed reduced efficiency of neural networks in older adults. The comparison of age groups for the semantic fluency task compared to rest revealed stronger activation only for the older adults. Significant clusters comprised hubs of the MDN as well as the right inferior frontal gyrus which has been emphasized in studies on semantic fluency in aging before (Meinzer et al., 2009; Meinzer et al., 2012a; Meinzer et al., 2012b; Nagels et al., 2012). Remarkably, the interaction between both tasks and age revealed significant effects only for the young adults who displayed stronger activation of frontal key regions of the MDN including the pre-SMA and dACC for semantic fluency. This finding suggests a pattern of increased processing efficiency which was reflected by faster response times compared to the older group. It converges with previous studies on language production and comprehension that associated greater activation in the prefrontal cortex with an increased task demand in young adults (Fu et al., 2002; Thompson-Schill et al., 1997; Whitney et al., 2009). The supplementary activation of MDN regions in older adults for the semantic fluency task compared to rest aligns with a meta-analysis on semantic cognition that found greater activity in areas of the MDN with older age (Hoffman and Morcom, 2018). The nature of this upregulation in brain activity in older adults has been subject of some debate (Cabeza et al., 2018; Morcom and Johnson, 2015). Here, we observed additional activation in the older adults while they performed poorer than the young adults during the more demanding semantic fluency task. In light of the additional beneficial activation of frontal MDN regions in the young adults, the observed upregulation in the older adults seems to further support the idea of age-related reduced efficiency of neural responses (Nyberg et al., 2014) leading to a stronger involvement of executive control at a lower level of task demand (Gallen et al, 2016; Hakun et al., 2015). This interpretation is backed up by the observed age-related differences in task-dependent activation of the MD and DM regions. During semantic fluency, older adults showed less deactivation of the DMN than young adults while during counting, the MDN was less deactivated in older than in young adults. Thus, in line with our functional connectivity results, older adults recruit similar neural resources as young adults, albeit at a lower level of processing efficiency which lends additional support to the hypothesis of dedifferentiation (Morcom and Henson, 2018; Park et al., 2004).

It should be noted that our results did not show a consistent effect of the intended modulation of task difficulty within the semantic fluency task on neural activation patterns. This could be related to the limited number of items participants had to produce for each category. A recent behavioral investigation on semantic fluency showed that the amount of correct responses continuously decreases with time (Gordon et al., 2018). Thus, although the effect of difficulty was present in the behavioral results in the form of reduced accuracy and slower responses, we assume that 9 trials per category were not enough to establish this effect on the neural level.

## Conclusion

In conclusion, the current study sheds light on the age-dependent contribution of the domain-general MD and DM systems during a verbal semantic fluency task. While univariate results revealed strong activity in the MDN during task processing, functional connectivity analyses demonstrated a strong interaction between the MDN and the DMN for semantic fluency. This finding corroborates the notion that usually anti-correlated networks integrate for successful task processing, especially when access to semantic memory is required. Although the strength of functional connectivity within- and between-networks was age-invariant, it had a different behavioral relevance in both age groups. Only the young adults engaged task-relevant networks in a beneficial way. This was evident in the form of better processing efficiency during semantic fluency and generally improved semantic memory. In older adults, strengthened functional connectivity within the MDN had a positive effect on cognitive performance, albeit older adults still performed at a lower level than young adults. Our results provide new insights into the concept of age-related reduced efficiency in the domain of semantic cognition and inform about the behavioral relevance of the frequently observed pattern of neural dedifferentiation.

## Acknowledgments

The authors would like to thank the medical technical assistants of MPI CBS for their support with data acquisition, and Annika Dunau, Caroline Duchow, and Rebekka Luckner for their support with transcriptions of recordings.

## Code and Data Availability

All behavioral data as well as extracted beta weights generated or analyzed during this study have been deposited in a public repository on Gitlab https://gitlab.gwdg.de/functionalconnectivityaging/mdn_lang. This repository also holds all self-written analysis code used for this project. Unthresholded statistical group maps for fMRI and gPPI results are made publicly available on NeuroVault: https://neurovault.org/collections/9072/. Raw and single subject neuroimaging data are protected under the General Data Protection Regulation (EU) and can only be made available from the authors upon reasonable request.

