## Supplementary Material for "Age-dependent contribution of domain-general networks to semantic cognition"

–

### Supplementary Material

#### Materials and Methods

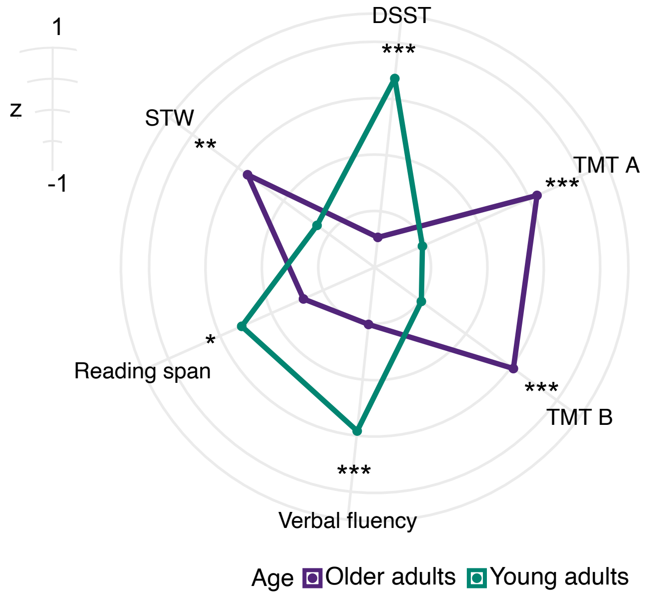


**Supplementary Figure S1. Age differences in neuropsychological tests.** STW = Spot-the-word test, DSST = Digit symbol substitution test, TMT A/B = Trail making test A/B. *** $p < 0.001$, ** $p < 0.01$, * $p < 0.05$.

#### Supplementary Results

##### Behavioral Result Tables

Regression tables were generated using RStudio (R Core Team, 2018) and the package sjPlot (Lüdecke, 2020).

**Table S1.** Results for mixed-effects models for accuracy and response time.

| | Accuracy | | | Response time | | |
|---|---|---|---|---|---|---|
| *Coefficient* | *Log-Odds* | *Conf. Int (95%)* | *p* | *Estimates* | *Conf. Int (95%)* | *p* |
| Intercept | 6.28 | 5.28 – 7.28 | **< 0.001** | 6.45 | 6.41 – 6.49 | **< 0.001** |
| Age | -3.82 | -5.66 – -1.97 | 0.662 | 0.02 | 0.01 – 0.03 | **< 0.001** |
| Condition | -6.47 | -8.44 – -4.50 | **< 0.001** | 0.17 | 0.13 – 0.22 | **< 0.001** |
| Difficulty | 4.63 | 2.67 – 6.59 | **< 0.001** | -0.06 | -0.11 – -0.01 | **< 0.001** |
| Age * Condition | 7.26 | 3.57 – 10.94 | 0.162 | 0.08 | 0.06 – 0.09 | **< 0.001** |
| Age * Difficulty | -7.99 | -11.64 – -4.34 | **0.002** | 0.00 | -0.01 – 0.02 | 0.7 |
| Condition * Difficulty | -5.03 | -8.94 – -1.11 | 0.049 | -0.10 | -0.19 – -0.00 | 0.056 |
| Age * Condition * Difficulty | 14.94 | 7.67 – 22.21 | **0.002** | 0.03 | -0.00 – 0.07 | 0.091 |
| **Random Effects** | | | | | | |
| $\sigma^2$ | 3.29 | | | 0.10 | | |
| $\tau_{00}$ | 0.19 $_{Subj}$ | | | 0.01 $_{Subj}$ | | |
| | 0.22 $_{Category}$ | | | 0.00 $_{Category}$ | | |
| ICC | 0.11 | | | 0.08 | | |
| N | 58 $_{Subj}$ | | | 58 $_{Subj}$ | | |
| | 22 $_{Category}$ | | | 22 $_{Category}$ | | |
| Observations | 19710 | | | 19491 | | |
| Marginal $R^2$ / Conditional $R^2$ | 0.900 / 0.911 | | | 0.079 / 0.156 | | |

Significant effects are marked in bold. Contrasts are sum coded. P-values were obtained via likelihood ratio tests. Conf. Int. Confidence interval.

**Table S2.** Results of post-hoc tests for three-way interaction Age x Condition x Difficulty for accuracy model. P-values are Bonferroni-corrected.

| **Contrast** | **Condition** | ***Odds ratio*** | ***SE*** | ***df*** | ***Conf. Int (95%)*** | ***z*** | ***p*** |
|---|---|---|---|---|---|---|---|
| OA Easy / YA Easy | Categories | 0.64 | 0.13 | Inf | 0.38 – 1.08 | -2.26 | 0.145 |
| OA Easy / OA Difficult | Categories | 6.41 | 1.69 | Inf | 3.20 – 12.83 | 7.06 | **< 0.001** |
| OA Easy / YA Difficult | Categories | 6.91 | 1.81 | Inf | 3.46 –13.80 | 7.36 | **< 0.001** |
| YA Easy / OA Difficult | Categories | 10.02 | 2.76 | Inf | 4.85 – 20.71 | 8.38 | **< 0.001** |
| YA Easy / YA Difficult | Categories | 10.79 | 2.95 | Inf | 5.25 – 22.21 | 8.7 | **< 0.001** |
| OA Difficult / YA Difficult | Categories | 1.08 | 0.09 | Inf | 0.86 – 1.35 | 0.86 | 1 |
| OA Easy / YA Easy | Counting | 0.00 | 0.00 | Inf | 0.00 – 0.00 | -4.1 | **< 0.001** |
| OA Easy / OA Difficult | Counting | 0.56 | 0.46 | Inf | 0.06 – 4.99 | -0.71 | 1 |
| OA Easy / YA Difficult | Counting | 0.74 | 0.59 | Inf | 0.09 – 6.12 | -0.38 | 1 |
| YA Easy / OA Difficult | Counting | 2167280.49 | 8196077.90 | Inf | 100.66 – 46660808797.59 | 3.86 | **< 0.001** |
| YA Easy / YA Difficult | Counting | 2884921.64 | 10821337.03 | Inf | 145.32 – 57273715280.44 | 3.97 | **< 0.001** |
| OA Difficult / YA Difficult | Counting | 1.33 | 0.69 | Inf | 0.34 – 5.21 | 0.55 | 1 |

Significant effects are marked in bold. SE standard error; df degrees of freedom; Conf. Int confidence interval.

**Table S3.** Results of post-hoc tests for three-way interaction Age x Condition x Difficulty for response time model. P-values are Bonferroni-corrected.

| Contrast | Condition | *Ratio* | *SE* | *df* | *Conf. Int (95%)* | *z* | *p* |
|---|---|---|---|---|---|---|---|
| OA Easy / YA Easy | Categories | 1.07 | 0.01 | Inf | 1.04 – 1.1 | 7.33 | **< 0.001** |
| OA Easy / OA Difficult | Categories | 0.91 | 0.02 | Inf | 0.87 – 0.95 | -5.69 | **< 0.001** |
| OA Easy / YA Difficult | Categories | 0.95 | 0.02 | Inf | 0.91 – 0.99 | -2.94 | **0.019** |
| YA Easy / OA Difficult | Categories | 0.85 | 0.01 | Inf | 0.81 – 0.89 | -9.60 | **< 0.001** |
| YA Easy / YA Difficult | Categories | 0.89 | 0.02 | Inf | 0.85 – 0.93 | -6.90 | **< 0.001** |
| OA Difficult / YA Difficult | Categories | 1.05 | 0.01 | Inf | 1.02 – 1.07 | 5.22 | **< 0.001** |
| OA Easy / YA Easy | Counting | 0.98 | 0.01 | Inf | 0.95 – 1 | -2.61 | 0.054 |
| OA Easy / OA Difficult | Counting | 0.98 | 0.05 | Inf | 0.87 – 1.11 | -0.32 | 1 |
| OA Easy / YA Difficult | Counting | 0.97 | 0.05 | Inf | 0.86 – 1.1 | -0.58 | 1 |
| YA Easy / OA Difficult | Counting | 1.01 | 0.05 | Inf | 0.89 – 1.14 | 0.18 | 1 |
| YA Easy / YA Difficult | Counting | 1 | 0.05 | Inf | 0.88 – 1.13 | -0.08 | 1 |
| OA Difficult / YA Difficult | Counting | 0.99 | 0.01 | Inf | 0.96 – 1.01 | -1.32 | 1 |

Significant effects are marked in bold. SE standard error; df degrees of freedom; Conf. Int confidence interval.

Functional MRI Results – within-group comparisons

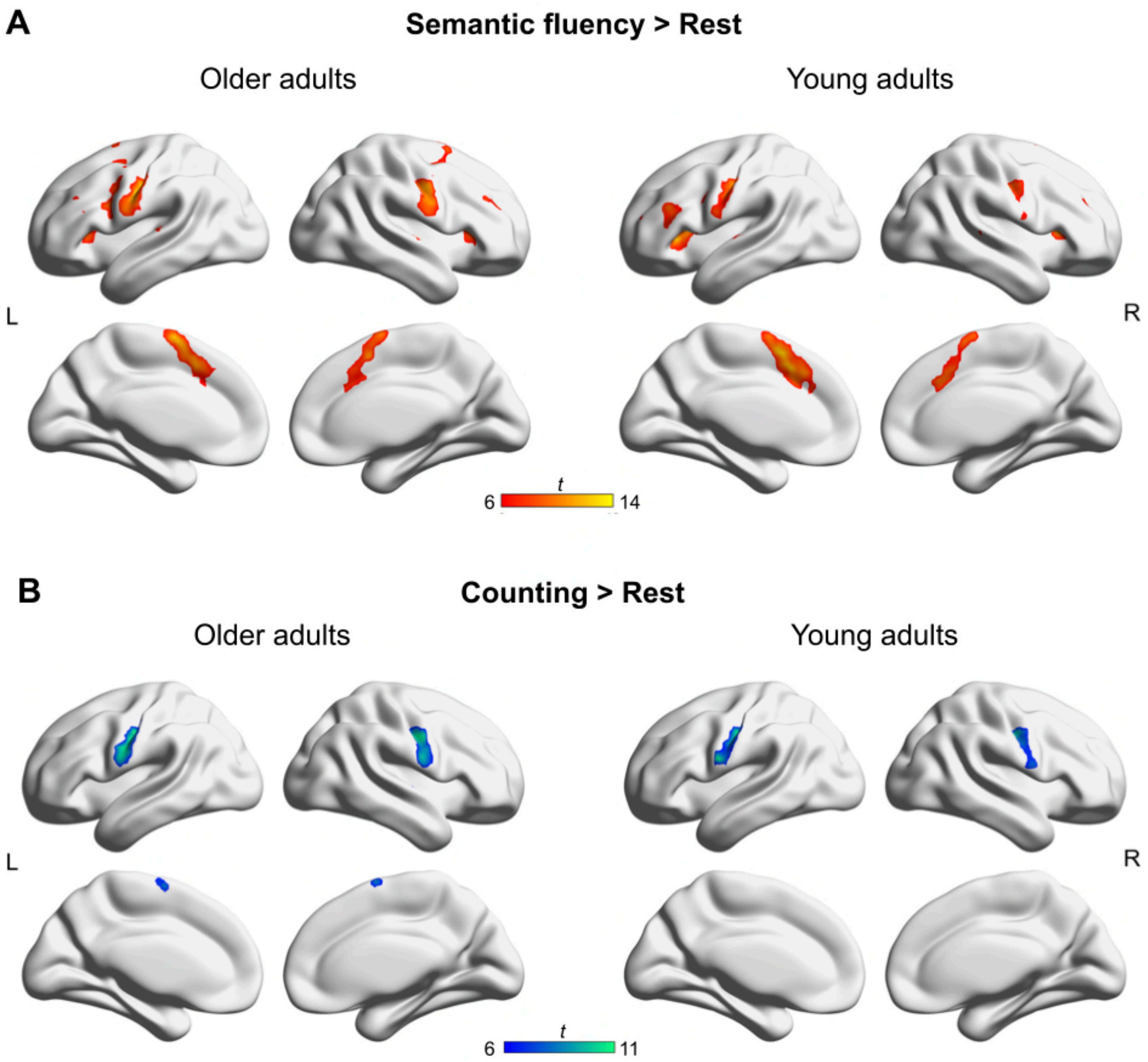


**Supplementary Figure S2. Functional MRI results for main effects of tasks from univariate analyses for each age group.** Results are FWE-corrected at $p < 0.05$ at peak-level with a minimum cluster size = 20 voxel. Unthresholded statistical maps are available at https://neurovault.org/collections/9072/.

**Functional MRI Activation Tables – within-group comparisons**

All X, Y, and Z coordinates are in Montreal Neurological Institute (MNI) atlas space. Cluster size (k) is given in $mm^3$.

**Table S4.** Older adults: Semantic fluency > Rest.

| Anatomical structure | Hemi | *k* | *t* | *x* | *y* | *z* |
|---|---|---|---|---|---|---|
| **Postcentral gyrus** | **L** | **899** | **14.26** | **-46** | **-10** | **35** |
| Postcentral gyrus | L | | 12.75 | -56 | -8 | 29 |
| Postcentral gyrus | L | | 12.01 | -51 | -13 | 46 |
| Postcentral gyrus | L | | 11.13 | -61 | 0 | 24 |
| **Cerebellum** | **L** | **407** | **13.28** | **-34** | **-57** | **-26** |
| Cerebellum | L | | 11.9 | -16 | -62 | -15 |
| Cerebellum | L | | 9.27 | -14 | -60 | -23 |
| Cerebellum | L | | 9 | -16 | -75 | -20 |
| **Supplementary motor cortex** | **L** | **878** | **13.11** | **-4** | **2** | **62** |
| Supplementary motor cortex | R | | 12.98 | 4 | 0 | 68 |
| Supplementary motor cortex | L | | 11.61 | -9 | 7 | 57 |
| Superior frontal gyrus | R | | 11.5 | 11 | 5 | 62 |
| **Postcentral gyrus** | **R** | **370** | **12** | **53** | **-5** | **26** |
| Precentral gyrus | R | | 11.77 | 51 | -5 | 35 |
| Postcentral gyrus | R | | 8.46 | 66 | -3 | 18 |
| **Caudate nucleus** | **R** | **86** | **11.63** | **18** | **2** | **24** |
| Caudate nucleus | R | | 9.59 | 16 | -8 | 21 |
| Caudate nucleus | R | | 8.72 | 11 | 0 | 10 |
| **Cerebellum** | **R** | **571** | **11.58** | **31** | **-65** | **-26** |
| Cerebellum | R | | 9.98 | 28 | -67 | -56 |
| Cerebellum | R | | 9.14 | 21 | -72 | -53 |
| Cerebellum | R | | 8.97 | 36 | -55 | -50 |
| **Middle frontal gyrus** | **R** | **163** | **11.36** | **36** | **47** | **26** |
| Middle frontal gyrus | R | | 9.23 | 41 | 35 | 29 |
| Middle frontal gyrus | R | | 7.43 | 26 | 35 | 26 |
| Middle frontal gyrus | R | | 7.28 | 36 | 42 | 18 |
| **Insula** | **L** | **122** | **9.92** | **-31** | **27** | **4** |
| Insula | L | | 8.09 | -31 | 15 | 7 |
| Inferior frontal gyrus, pars orbitalis | L | | 7.92 | -44 | 37 | -9 |
| **Insula** | **R** | **148** | **9.91** | **31** | **27** | **2** |
| Insula | R | | 8.77 | 43 | 20 | 2 |
| Inferior frontal gyrus, pars orbitalis | R | | 6.47 | 38 | 32 | -6 |
| **Superior temporal gyrus** | **R** | **78** | **9.76** | **63** | **-3** | **2** |
| Superior temporal gyrus | R | | 8.24 | 56 | -13 | 2 |
| Superior temporal gyrus | R | | 7.49 | 68 | -15 | 2 |

| | | | | | | |
|---|---|---|---|---|---|---|
| Superior temporal gyrus | R | | 7.06 | 51 | -18 | 7 |
| **Caudate nucleus** | **L** | **45** | **9.56** | **-16** | **-10** | **24** |
| Caudate nucleus | L | | 9.11 | -14 | -5 | 16 |
| **Superior parietal lobe** | **L** | **98** | **9.33** | **-19** | **-60** | **48** |
| Inferior parietal sulcus | L | | 8.5 | -26 | -45 | 38 |
| Inferior parietal lobe | L | | 7.99 | -26 | -55 | 38 |
| Inferior parietal lobe | L | | 7.77 | -41 | -40 | 38 |
| **Superior temporal gyrus** | **L** | **54** | **8.72** | **-51** | **-28** | **10** |
| Superior temporal gyrus | L | | 7.68 | -66 | -23 | 10 |
| Superior temporal gyrus | L | | 6.7 | -41 | -32 | 10 |
| **Inferior frontal gyrus, pars triangularis** | **L** | **20** | **8.71** | **-36** | **40** | **4** |
| **Superior temporal gyrus** | **L** | **21** | **8.09** | **-64** | **-10** | **4** |

FWE-corrected ($p < 0.05$) at peak level, $k \geq 20$ voxels.

**Table S5.** Young adults: Semantic fluency > Rest.

| Anatomical structure | Hemi | *k* | *t* | *x* | *y* | *z* |
|---|---|---|---|---|---|---|
| **Presupplementary motor cortex** | **L** | **767** | **15.05** | **-4** | **12** | **51** |
| Supplementary motor cortex | L | | 13.89 | -6 | 17 | 43 |
| Supplementary motor cortex | R | | 12.54 | 4 | 7 | 62 |
| Middle cingulate cortex | R | | 11.17 | 11 | 20 | 38 |
| **Insula** | **L** | **280** | **13.95** | **-34** | **27** | **2** |
| Insula | L | | 11.74 | -31 | 20 | 7 |
| Inferior frontal gyrus, pars opercularis | L | | 8.61 | -46 | 10 | 7 |
| Inferior frontal gyrus, pars opercularis | L | | 7.01 | -49 | 17 | -4 |
| **Cerebellum** | **R** | **683** | **13.82** | **33** | **-55** | **-31** |
| Cerebellum | R | | 13.01 | 43 | -60 | -28 |
| Cerebellum | R | | 11.51 | 33 | -62 | -50 |
| Cerebellum | R | | 11.24 | 26 | -67 | -48 |
| **Postcentral gyrus** | **L** | **331** | **13.74** | **-49** | **-15** | **40** |
| Postcentral gyrus | L | | 10.89 | -56 | -13 | 46 |
| Postcentral gyrus | L | | 9.93 | -61 | 0 | 21 |
| Precentral gyrus | L | | 8.04 | -54 | 0 | 46 |
| **Cerebellum** | **L** | **377** | **13.18** | **-26** | **-60** | **-26** |
| Cerebellum | L | | 12.1 | -46 | -62 | -28 |
| **Insula** | **R** | **130** | **12.01** | **33** | **22** | **7** |
| Inferior frontal gyrus, pars opercularis | R | | 9.4 | 46 | 15 | 4 |
| Insula | R | | 9.05 | 41 | 20 | -1 |
| **Cerebellum** | **R** | **40** | **10.86** | **1** | **-47** | **-23** |

| | | | | | | |
|---|---|---|---|---|---|---|
| **Cerebellum** | **L** | **68** | **10.41** | **-36** | **-60** | **-50** |
| **Precentral gyrus** | **R** | **208** | **10.23** | **56** | **-3** | **46** |
| Precentral gyrus | R | | 10.09 | 46 | -10 | 38 |
| Postcentral gyrus | R | | 8.94 | 56 | -5 | 35 |
| Rolandic operculum | R | | 7.71 | 61 | -3 | 16 |
| **Inferior frontal gyrus, pars triangularis** | **L** | **208** | **9.99** | **-44** | **32** | **24** |
| Inferior frontal gyrus, pars triangularis | L | | 8.29 | -51 | 30 | 21 |
| Inferior frontal gyrus, pars triangularis | L | | 8.13 | -39 | 35 | 7 |
| Inferior frontal gyrus, pars triangularis | L | | 7.96 | -51 | 35 | 10 |
| **Thalamus** | **L** | **57** | **9.31** | **-11** | **-5** | **13** |
| Caudate nucleus | L | | 9.2 | -16 | -3 | 21 |
| **Inferior frontal gyrus, pars opercularis** | **L** | **98** | **8.33** | **-39** | **2** | **26** |
| Precentral gyrus | L | | 7.6 | -46 | 10 | 32 |
| Precentral gyrus | L | | 7.16 | -41 | 0 | 38 |
| Inferior frontal gyrus, pars triangularis | L | | 6.5 | -46 | 15 | 24 |
| **Middle frontal gyrus** | **R** | **81** | **8.1** | **33** | **47** | **32** |
| Middle frontal gyrus | R | | 7.08 | 31 | 47 | 24 |
| **Caudate nucleus** | **R** | **43** | **8.09** | **18** | **5** | **21** |
| Caudate nucleus | R | | 7.75 | 16 | -3 | 24 |
| Caudate nucleus | R | | 7.73 | 18 | 12 | 16 |
| **Middle frontal gyrus** | **L** | **26** | **7.57** | **-34** | **55** | **21** |
| **Superior temporal gyrus** | **R** | **21** | **7.5** | **66** | **-30** | **7** |
| Superior temporal gyrus | R | | 6.7 | 56 | -30 | 4 |
| **Superior temporal gyrus** | **L** | **20** | **6.74** | **-59** | **-15** | **4** |

FWE-corrected ($p < 0.05$) at peak level, $k \geq 20$ voxels.

**Table S6.** Older adults: Counting > Rest.

| Anatomical structure | Hemi | *k* | *t* | *x* | *y* | *z* |
|---|---|---|---|---|---|---|
| **Postcentral gyrus** | **L** | **347** | **11.69** | **-46** | **-13** | **38** |
| Postcentral gyrus | L | | 11.56 | -61 | -3 | 24 |
| Postcentral gyrus | L | | 11.52 | -51 | -13 | 46 |
| Postcentral gyrus | L | | 11.05 | -56 | -8 | 29 |
| **Postcentral gyrus** | **R** | **342** | **11.43** | **48** | **-8** | **35** |
| Postcentral gyrus | R | | 10.71 | 53 | -3 | 24 |
| Postcentral gyrus | R | | 9.39 | 63 | -3 | 18 |
| **Supplementary motor cortex** | **L** | **57** | **10.16** | **-4** | **-5** | **70** |
| **Supplementary motor cortex** | **R** | **59** | **9.96** | **4** | **0** | **68** |

| | | | | | | |
|---|---|---|---|---|---|---|
| **Cerebellum** | **L** | **52** | **9.35** | **-29** | **-62** | **-23** |
| Cerebellum | L | | 7.84 | -16 | -65 | -18 |
| **Superior temporal gyrus** | **R** | **49** | **8.44** | **66** | **-10** | **2** |
| Superior temporal gyrus | R | | 7.71 | 63 | 2 | -1 |
| Superior temporal gyrus | R | | 6.96 | 68 | -18 | 4 |
| Superior temporal gyrus | R | | 6.76 | 53 | -15 | 4 |
| **Superior temporal gyrus** | **L** | **46** | **8.4** | **-46** | **-42** | **21** |
| Superior temporal gyrus | L | | 6.54 | -46 | -37 | 13 |
| Superior temporal gyrus | L | | 6.19 | -51 | -28 | 10 |
| **Cerebellum** | **R** | **26** | **6.96** | **21** | **-60** | **-23** |
| Cerebellum | R | | 6.9 | 11 | -60 | -23 |

FWE-corrected (p < 0.05) at peak level, k ≥ 20 voxels.

**Table S7.** Young adults: Counting > Rest.

| Anatomical structure | Hemi | *k* | *t* | *x* | *y* | *z* |
|---|---|---|---|---|---|---|
| **Postcentral gyrus** | **L** | **317** | **11.47** | **-61** | **0** | **21** |
| Postcentral gyrus | L | | 11.3 | -49 | -15 | 40 |
| Postcentral gyrus | L | | 9.28 | -56 | -13 | 46 |
| Postcentral gyrus | L | | 6.99 | -59 | -8 | 16 |
| **Precentral gyrus** | **R** | **247** | **10.42** | **46** | **-10** | **38** |
| Precentral gyrus | R | | 9.44 | 56 | -3 | 46 |
| Rolandic operculum | R | | 9.35 | 61 | 2 | 16 |
| Postcentral gyrus | R | | 8.25 | 56 | -8 | 35 |
| **Cerebellum** | **R** | **37** | **8.3** | **13** | **-60** | **-20** |
| **Cerebellum** | **L** | **25** | **7.37** | **-16** | **-60** | **-23** |

FWE-corrected (p < 0.05) at peak level, k ≥ 20 voxels.

**Table S8.** Older adults: Semantic fluency > Counting.

| Anatomical structure | Hemi | *k* | *t* | *x* | *y* | *z* |
|---|---|---|---|---|---|---|
| **Cerebellum** | **L** | **1844** | **12.99** | **-36** | **-62** | **-26** |
| Cerebellum | R | | 12.59 | 28 | -62 | -26 |
| Cerebellum | L | | 12.03 | -29 | -67 | -26 |
| Cerebellum | R | | 11.08 | 6 | -80 | -31 |
| **Middle frontal gyrus** | **L** | **578** | **12.9** | **-44** | **5** | **35** |
| Inferior frontal gyrus, pars opercularis | L | | 11.2 | -41 | 15 | 21 |
| Precentral gyrus | L | | 10.7 | -39 | 2 | 24 |
| Middle frontal gyrus | L | | 8.94 | -46 | 7 | 46 |
| **Superior frontal gyrus (preSMA)** | **L** | **661** | **12.02** | **-9** | **15** | **51** |
| Presupplementary motor cortex | L | | 11.62 | -9 | 20 | 43 |
| Superior frontal gyrus | L | | 10.34 | -1 | 10 | 60 |

| | | | | | | |
|---|---|---|---|---|---|---|
| Superior frontal gyrus | R | | 9.34 | 8 | 15 | 48 |
| **Insula** | **L** | **269** | **10.98** | **-31** | **25** | **4** |
| Caudate nucleus | L | | 10.84 | -16 | 0 | 16 |
| Caudate nucleus | L | | 9.52 | -16 | -10 | 21 |
| Superior frontal gyrus | L | | 8.74 | -19 | 10 | 4 |
| **Insula** | **R** | **113** | **10.95** | **31** | **27** | **2** |
| Inferior frontal gyrus. pars triangularis | R | | 7.25 | 48 | 22 | -4 |
| Frontal operculum | R | | 7.08 | 43 | 20 | 4 |
| **Caudate nucleus** | **R** | **106** | **9.96** | **18** | **15** | **18** |
| Caudate nucleus | R | | 9.53 | 18 | -8 | 21 |
| Caudate nucleus | R | | 8.67 | 16 | 0 | 18 |
| Thalamus | R | | 8.22 | 11 | 0 | 10 |
| **Middle frontal gyrus** | **R** | **36** | **9.79** | **43** | **35** | **32** |
| **Superior frontal gyrus** | **L** | **100** | **9.12** | **-21** | **12** | **54** |
| Middle frontal gyrus | L | | 6.82 | -21 | -3 | 60 |
| Middle frontal gyrus | L | | 6.57 | -24 | 20 | 51 |
| **Intracalcarine cortex** | **R** | **43** | **9.01** | **18** | **-80** | **7** |
| Occipital pole | R | | 7.93 | 13 | -95 | 10 |
| Intracalcarine cortex | R | | 6.77 | 13 | -77 | 16 |
| **Angular gyrus** | **L** | **27** | **8.1** | **-34** | **-72** | **43** |
| **Middle frontal gyrus** | **L** | **24** | **8.07** | **-34** | **0** | **57** |
| **Superior parietal lobe** | **L** | **61** | **7.65** | **-14** | **-65** | **51** |
| Superior parietal lobe | L | | 7.31 | -21 | -65 | 60 |
| Angular gyrus | L | | 6.65 | -29 | -62 | 46 |
| **Intracalcarine cortex** | **L** | **71** | **7.52** | **-11** | **-72** | **10** |
| Intracalcarine cortex | L | | 7.24 | -6 | -87 | 2 |
| Intracalcarine cortex | L | | 6.96 | -4 | -82 | 10 |
| **Middle frontal gyrus** | **R** | **21** | **7.39** | **23** | **60** | **-4** |
| Middle frontal gyrus | R | | 7.05 | 31 | 57 | -9 |
| **Thalamus** | **L** | **21** | **6.99** | **-4** | **-5** | **10** |

FWE-corrected ($p < 0.05$) at peak level, $k \geq 20$ voxels.

**Table S9.** Young adults: Semantic fluency > Counting.

| **Anatomical structure** | **Hemi** | ***k*** | ***t*** | ***x*** | ***y*** | ***z*** |
|---|---|---|---|---|---|---|
| **Insula** | **L** | **3112** | **20.03** | **-31** | **25** | **2** |
| Presupplementary motor cortex | L | | 16.85 | -4 | 25 | 40 |
| Presupplementary motor cortex | L | | 16.33 | -6 | 12 | 51 |
| Presupplementary motor cortex | R | | 14.72 | 13 | 27 | 29 |
| **Cerebellum** | **R** | **2970** | **19.11** | **33** | **-57** | **-31** |
| Cerebellum | R | | 16.12 | 31 | -65 | -28 |
| Cerebellum | R | | 13.89 | 28 | -70 | -50 |
| Cerebellum | R | | 12.66 | 41 | -60 | -28 |

| | | | | | | |
|---|---|---|---|---|---|---|
| **Anterior cingulate gyrus** | **L** | **90** | **14.15** | **-4** | **2** | **29** |
| Anterior cingulate gyrus | L | | 9.12 | -1 | 12 | 24 |
| Anterior cingulate gyrus | R | | 8.89 | 6 | 7 | 26 |
| **Insula** | **R** | **315** | **13.32** | **31** | **27** | **2** |
| Insula | R | | 12.64 | 38 | 20 | -4 |
| **Caudate nucleus** | **L** | **235** | **12.81** | **-9** | **5** | **2** |
| Caudate nucleus | L | | 12.02 | -16 | -3 | 21 |
| Thalamus | L | | 11.32 | -11 | -5 | 13 |
| Caudate nucleus | L | | 10.63 | -16 | 7 | 16 |
| **Brain stem** | **L** | **296** | **12.46** | **-6** | **-23** | **-18** |
| Thalamus | L | | 11.94 | -9 | -18 | 16 |
| Thalamus | L | | 11.85 | -4 | -23 | 10 |
| Thalamus | L | | 10.36 | -4 | -13 | 10 |
| **Superior parietal lobe** | **L** | **224** | **10.49** | **-29** | **-65** | **51** |
| Angular gyrus | L | | 10.21 | -29 | -72 | 43 |
| Inferior parietal lobe | L | | 8.84 | -34 | -57 | 40 |
| Middle occipital gyrus | L | | 6.47 | -31 | -80 | 38 |
| **Caudate nucleus** | **R** | **215** | **10.45** | **18** | **10** | **18** |
| Caudate nucleus | R | | 10.38 | 8 | 7 | 2 |
| Caudate nucleus | R | | 9.57 | 13 | 7 | 10 |
| Caudate nucleus | R | | 9.16 | 18 | -3 | 21 |
| **Superior temporal gyrus** | **L** | **54** | **9.62** | **-61** | **-30** | **7** |
| Planum temporale | L | | 6.97 | -61 | -15 | 4 |
| **Middle frontal gyrus** | **R** | **176** | **8.3** | **36** | **42** | **32** |
| Middle frontal gyrus | R | | 7.56 | 31 | 55 | 26 |
| Middle frontal gyrus | R | | 7.53 | 33 | 37 | 21 |
| Middle frontal gyrus | R | | 7.17 | 41 | 35 | 40 |

FWE-corrected ($p < 0.05$) at peak level, $k \geq 20$ voxels.

**Table S10.** Older adults: Counting > Semantic fluency.

| Anatomical structure | Hemi | *k* | *t* | *x* | *y* | *z* |
|---|---|---|---|---|---|---|
| **Temporal pole** | **R** | **30** | **9.33** | **51** | **12** | **-31** |
| **Precuneus** | **R** | **45** | **7.72** | **6** | **-52** | **38** |

FWE-corrected ($p < 0.05$) at peak level, $k \geq 20$ voxels.

**Table S11.** Young adults: Counting > Semantic fluency (FWE-corrected).

| Anatomical structure | Hemi | *k* | *t* | *x* | *y* | *z* |
|---|---|---|---|---|---|---|
| **Temporal pole** | **R** | **75** | **11.02** | **51** | **10** | **-31** |
| Temporal pole | R | | 6.93 | 43 | 20 | -28 |
| **Precuneus** | **R** | **312** | **9.7** | **8** | **-65** | **29** |
| Precuneus | R | | 9.59 | 11 | -52 | 35 |
| Precuneus | L | | 9.05 | -9 | -52 | 35 |

| | | | | | | |
|---|---|---|---|---|---|---|
| **Insula** | **L** | **46** | **8.62** | **-41** | **-8** | **-1** |
| Insula | L | | 7.24 | -36 | -18 | 18 |
| Insula | L | | 7.02 | -39 | -15 | 2 |
| **Insula** | **R** | **62** | **8.48** | **36** | **-15** | **4** |
| Insula | R | | 7.66 | 41 | 0 | -6 |
| Insula | R | | 7.1 | 38 | -15 | 21 |
| **Middle temporal gyrus** | **L** | **27** | **8.27** | **-56** | **2** | **-20** |
| **Rolandic operculum** | **R** | **22** | **8.01** | **53** | **0** | **10** |
| **Posterior cingulate cortex** | **L** | **40** | **7.9** | **-6** | **-30** | **46** |
| Precentral gyrus | L | | 6.77 | -6 | -25 | 54 |
| **Precentral gyrus** | **R** | **28** | **7.42** | **1** | **-15** | **62** |
| Precentral gyrus | L | | 6.91 | -4 | -23 | 70 |

FWE-corrected ($p < 0.05$) at peak level, $k \geq 20$ voxels.

**Table S12.** Young adults: Counting > Semantic fluency ($p < 0.001$ uncorr., FWE-corrected $p < 0.05$ at cluster level).

| Anatomical structure | Hemi | *k* | *t* | *x* | *y* | *z* |
|---|---|---|---|---|---|---|
| **Temporal pole** | **R** | **281** | **11.02** | **51** | **10** | **-31** |
| Temporal pole | R | | 6.93 | 43 | 20 | -28 |
| Middle temporal gyrus | R | | 5.93 | 61 | -8 | -12 |
| Superior temporal gyrus | R | | 4.35 | 48 | -10 | -15 |
| **Precuneus** | **R** | **3620** | **9.7** | **8** | **-65** | **29** |
| Precuneus | R | | 9.59 | 11 | -52 | 35 |
| Precuneus | L | | 9.05 | -9 | -52 | 35 |
| Insula | R | | 8.48 | 36 | -15 | 4 |
| Central operculum | R | | 8.01 | 53 | 0 | 10 |
| **Insula** | **L** | **438** | **8.62** | **-41** | **-8** | **-1** |
| Insula | L | | 7.24 | -36 | -18 | 18 |
| Insula | L | | 7.02 | -39 | -15 | 2 |
| Precentral gyrus | L | | 6.99 | -59 | 2 | 10 |
| Insula | L | | 5.22 | -41 | -3 | -12 |
| **Middle temporal gyrus** | **L** | **151** | **8.27** | **-56** | **2** | **-20** |
| Temporal pole | L | | 7.06 | -54 | 10 | -31 |
| Temporal pole | L | | 4.5 | -44 | 20 | -31 |
| Temporal pole | L | | 3.99 | -41 | 7 | -23 |
| **Anterior cingulate cortex** | **L** | **96** | **7.36** | **-6** | **27** | **-6** |
| Anterior cingulate cortex | L | | 4.09 | -6 | 42 | -4 |
| **Angular gyrus** | **L** | **281** | **7.11** | **-54** | **-62** | **35** |
| Angular gyrus | L | | 5.46 | -41 | -60 | 26 |
| Middle temporal gyrus | L | | 5.05 | -46 | -62 | 18 |
| Angular gyrus | L | | 4.46 | -46 | -75 | 35 |
| Angular gyrus | L | | 4.03 | -49 | -67 | 43 |
| **Angular gyrus** | **R** | **389** | **7.02** | **51** | **-57** | **26** |

| | | | | | |
|---|---|---|---|---|---|
| Angular gyrus | R | 4.6 | 46 | -65 | 48 |
| Angular gyrus | R | 4.16 | 43 | -72 | 35 |
| Lateral occipital cortex | R | 4.1 | 46 | -77 | 26 |
| Lateral occipital cortex | R | 3.55 | 56 | -62 | 7 |

FWE-corrected ($p < 0.05$) at cluster-level, $p < 0.001$ uncorr. at peak-level.

**Table S13.** Results for linear mixed-effects model for parameter estimates from fMRI main effects.

| | Beta weights | | |
|---|---|---|---|
| *Coefficient* | *Estimates* | *Conf. Int (95%)* | *p* |
| Intercept | -0.39 | -0.54 – -0.24 | **< 0.001** |
| Network | 1.07 | 0.95 – 1.19 | **< 0.001** |
| Age | 0.34 | 0.21 – 0.46 | **< 0.001** |
| Condition | 0.49 | 0.37 – 0.61 | **< 0.001** |
| Network * Age | -0.02 | -0.15 – 0.10 | 0.704 |
| Network * Condition | 1.14 | 1.02 – 1.26 | **< 0.001** |
| Age * Condition | -0.02 | -0.14 – 0.10 | 0.762 |
| Network * Age * Condition | -0.27 | -0.39 – -0.15 | **< 0.001** |
| **Random Effects** | | | |
| $\sigma^2$ | 0.91 | | |
| $\tau_{00\ Subj}$ | 0.06 | | |
| ICC | 0.06 | | |
| $N_{Subj}$ | 30 | | |
| Observations | 232 | | |
| Marginal $R^2$ / Conditional $R^2$ | 0.751 / 0.766 | | |

Significant effects are marked in bold. Contrasts are sum coded. P-values were obtained via likelihood ratio tests. Conf. Int. Confidence interval.

Functional MRI Results – within-group comparisons

**Table S14.** Results for post-hoc tests for three-way interaction Network x Age x Contrast for parameter estimates model. P-values are Bonferroni-corrected.

| **Contrast** | **fMRI contrast** | ***Estimate*** | ***SE*** | ***df*** | ***Conf. Int (95%)*** | ***t*** | ***p*** |
|---|---|---|---|---|---|---|---|
| OA MDN - YA MDN | SF > rest | 0.04 | 0.25 | 196.93 | -0.63 – 0.71 | 0.17 | 1 |
| OA MDN - OA DMN | SF > rest | 3.83 | 0.25 | 195.3 | 3.15 – 4.51 | 15.05 | **< 0.001** |
| OA MDN - YA DMN | SF > rest | 5.05 | 0.25 | 196.93 | 4.38 – 5.72 | 20.18 | **< 0.001** |
| YA MDN - OA DMN | SF > rest | 3.79 | 0.25 | 196.93 | 3.12 – 4.45 | 15.12 | **< 0.001** |
| YA MDN - YA DMN | SF > rest | 5.01 | 0.25 | 195.3 | 4.35 – 5.66 | 20.38 | **< 0.001** |
| OA DMN - YA DMN | SF > rest | 1.22 | 0.25 | 196.93 | 0.56 – 1.89 | 4.89 | **< 0.001** |
| OA MDN - YA MDN | Count > rest | 1.20 | 0.25 | 196.93 | 0.54 – 1.87 | 4.81 | **< 0.001** |
| OA MDN - OA DMN | Count > rest | 0.35 | 0.25 | 195.3 | -0.33 – 1.03 | 1.38 | 1 |
| OA MDN - YA DMN | Count > rest | 0.57 | 0.25 | 196.93 | -0.10 – 1.24 | 2.27 | 0.146 |
| YA MDN - OA DMN | Count > rest | -0.85 | 0.25 | 196.93 | -1.52 – -0.19 | -3.41 | **0.005** |
| YA MDN - YA DMN | Count > rest | -0.64 | 0.25 | 195.3 | -1.29 – 0.02 | -2.59 | 0.062 |
| OA DMN - YA DMN | Count > rest | 0.22 | 0.25 | 196.93 | -0.45 – 0.89 | 0.87 | 1 |

Significant effects are marked in bold. SE standard error; df degrees of freedom; Conf. Int confidence interval.

**Table S 15.** Young adults: Easy > Difficult semantic categories.

| **Anatomical structure** | **Hemi** | ***k*** | ***t*** | ***x*** | ***y*** | ***z*** |
|---|---|---|---|---|---|---|
| **Middle frontal gyrus** | **R** | **20** | **7.24** | **26** | **2** | **51** |
| Middle frontal gyrus | R | | 6.49 | 31 | 12 | 51 |

FWE-corrected ($p < 0.05$) at peak level, $k \geq 20$ voxels.

**Activation Tables for Psychophysiological Interactions (PPI)**

All individual seeds for PPIs were thresholded at $p < 0.01$. All reported results are for the contrast Semantic fluency > Counting and are FWE-corrected at $p < 0.05$ at peak-level (k ≥ 20 voxels). All X, Y, and Z coordinates are in Montreal Neurological Institute (MNI) atlas space. Cluster size (k) is given in $mm^3$.

**Table S16.** PPI seed: Pre-supplementary Motor Area [-6 12 51].

| Anatomical structure | Hemi | *k* | *t* | *x* | *y* | *z* |
|---|---|---|---|---|---|---|
| **Older adults** | | | | | | |
| No significant clusters above threshold. | | | | | | |
| **Young adults** | | | | | | |
| **Caudate nucleus** | **L** | **92** | **9.89** | **-14** | **10** | **4** |
| Caudate nucleus | L | | 8.68 | -16 | 20 | 4 |
| Caudate nucleus | L | | 6.18 | -11 | 7 | 13 |
| **Caudate nucleus** | **R** | **76** | **9.19** | **8** | **12** | **2** |
| Caudate nucleus | R | | 7.72 | 18 | 22 | -4 |
| Putamen | R | | 7.30 | 18 | 12 | -1 |
| Caudate nucleus | R | | 6.93 | 18 | 25 | 4 |
| **Precuneus** | **L** | **125** | **8.31** | **-6** | **-52** | **16** |
| Posterior cingulate cortex | L | | 7.60 | -4 | -55 | 26 |
| Precuneus | L | | 7.44 | -11 | -55 | 7 |
| **Thalamus** | **L** | **35** | **7.30** | **-1** | **-13** | **7** |
| Thalamus | R | | 7.29 | 4 | -20 | 10 |
| Thalamus | L | | 6.50 | -9 | -25 | 13 |

**Table S17.** PPI seed: Left Insula [-31 25 2].

| Anatomical structure | Hemi | *k* | *t* | x | y | z |
|---|---|---|---|---|---|---|
| **Older adults** | | | | | | |
| No significant clusters above threshold. | | | | | | |
| **Young adults** | | | | | | |
| **Caudate nucleus** | **R** | **36** | **10.09** | **8** | **12** | **2** |
| **Caudate nucleus** | **L** | **34** | **7.04** | **-14** | **12** | **4** |
| Caudate nucleus | L | | 6.33 | -16 | 22 | 2 |
| **Precuneus** | **L** | **31** | **6.88** | **-9** | **-60** | **7** |
| Precuneus | L | | 6.75 | -6 | -52 | 16 |

**Table S18.** PPI seed: Right Insula [31 27 2].

| Anatomical structure | Hemi | *k* | *t* | *x* | *y* | *z* |
|---|---|---|---|---|---|---|
| **Older adults** | | | | | | |

| | | | | | | |
|---|---|---|---|---|---|---|
| **Precuneus** | **R** | **29** | **8.92** | **1** | **-60** | **26** |
| Posterior cingulate gyrus | L | | 8.56 | -1 | -45 | 26 |
| **Inferior frontal gyrus, pars orbitalis** | **L** | **20** | **7.39** | **-31** | **35** | **-15** |
| Inferior frontal gyrus, pars orbitalis | L | | 6.46 | -44 | 30 | -12 |
| **Young adults** | | | | | | |
| No significant clusters above threshold. | | | | | | |

**Table S19.** PPI seed: Right Temporal Pole [48 15 -31].

| Anatomical structure | Hemi | *k* | *t* | *x* | *y* | *z* |
|---|---|---|---|---|---|---|
| **Older adults** | | | | | | |
| **Inferior frontal gyrus, pars opercularis** | **R** | **24** | **7.33** | **51** | **17** | **-1** |
| Insula | R | | 7.02 | 41 | 27 | -1 |
| Insula | R | | 6.31 | 41 | 12 | -1 |
| **Young adults** | | | | | | |
| **Inferior frontal gyrus, pars opercularis** | **R** | **47** | **8.48** | **43** | **12** | **21** |
| Inferior frontal gyrus, pars opercularis | R | | 7.08 | 53 | 12 | 18 |
| **Superior frontal gyrus** | **R** | **46** | **8.28** | **18** | **2** | **65** |
| Middle frontal gyrus | R | | 6.94 | 31 | -3 | 62 |
| Superior frontal gyrus | R | | 6.44 | 21 | 12 | 65 |
| **Insula** | **R** | **75** | **7.22** | **43** | **15** | **2** |
| Frontal operculum | R | | 6.76 | 33 | 25 | 7 |
| Frontal operculum | R | | 6.53 | 36 | 15 | 10 |
| **Supramarginal gyrus** | **R** | **23** | **7.21** | **58** | **-32** | **48** |

**Table S20.** PPI seed: Right Precuneus [8 -65 29].

| Anatomical structure | Hemi | *k* | *t* | *x* | *y* | *z* |
|---|---|---|---|---|---|---|
| **Older adults** | | | | | | |
| **Insula** | **R** | **295** | **9.27** | **33** | **22** | **10** |
| Inferior frontal gyrus, pars triangularis | R | | 9.27 | 53 | 25 | 10 |
| Insula | R | | 8.97 | 41 | 25 | -6 |
| Inferior frontal gyrus, pars triangularis | R | | 8.79 | 46 | 25 | 4 |
| **Supramarginal gyrus** | **R** | **410** | **9.20** | **53** | **-40** | **46** |
| Angular gyrus | R | | 9.18 | 56 | -45 | 32 |
| Supramarginal gyrus | R | | 8.91 | 63 | -42 | 35 |
| Angular gyrus | R | | 8.72 | 56 | -47 | 48 |
| **Middle frontal gyrus** | **R** | **125** | **8.78** | **33** | **47** | **32** |
| Middle frontal gyrus | R | | 8.04 | 41 | 42 | 29 |
| Middle frontal gyrus | R | | 7.39 | 46 | 45 | 21 |
| **Supramarginal gyrus** | **L** | **42** | **8.77** | **-61** | **-47** | **32** |

| | | | | | | |
|---|---|---|---|---|---|---|
| Supramarginal gyrus | L | | 6.89 | -54 | -50 | 35 |
| **Inferior frontal gyrus, pars orbitalis** | **R** | **61** | **8.4** | **48** | **45** | **-6** |
| Inferior frontal gyrus, pars triangularis | R | | 8.03 | 51 | 37 | -1 |
| Inferior frontal gyrus, pars triangularis | R | | 7.78 | 48 | 40 | 7 |
| Inferior frontal gyrus, pars triangularis | R | | 7.38 | 41 | 42 | -1 |
| **Presupplementary motor area** | **R** | **35** | **8.06** | **4** | **7** | **60** |
| Presupplementary motor area | R | | 7.44 | 6 | 10 | 68 |
| **Inferior frontal gyrus, pars triangularis** | **L** | **70** | **8.02** | **-41** | **17** | **7** |
| Inferior frontal gyrus, pars triangularis | L | | 7.85 | -36 | 30 | 4 |
| Inferior frontal gyrus, pars opercularis | L | | 6.84 | -46 | 10 | 7 |
| **Superior temporal gyrus** | **R** | **32** | **7.86** | **53** | **-15** | **-4** |
| Middle temporal gyrus | R | | 7.03 | 56 | -30 | -1 |
| Superior temporal gyrus | R | | 6.77 | 63 | -20 | -1 |
| **Precentral gyrus** | **R** | **93** | **7.85** | **46** | **7** | **35** |
| Middle frontal gyrus | R | | 6.89 | 41 | 10 | 46 |
| Middle frontal gyrus | R | | 6.88 | 41 | 15 | 29 |
| Precentral gyrus | R | | 6.82 | 43 | 5 | 26 |
| **Inferior frontal gyrus, pars opercularis** | **R** | **21** | **7.77** | **56** | **15** | **24** |
| **Angular gyrus** | **L** | **39** | **7.61** | **-51** | **-52** | **48** |
| Supramarginal gyrus | L | | 7.19 | -59 | -47 | 43 |
| **Lateral occipital cortex** | **R** | **29** | **7.15** | **33** | **-67** | **29** |
| Lateral occipital cortex | R | | 6.78 | 26 | -77 | 26 |
| **Young adults** | | | | | | |
| **Supramarginal gyrus** | **R** | **1300** | **9.69** | **61** | **-45** | **26** |
| Angular gyrus | R | | 8.29 | 63 | -47 | 18 |
| Supramarginal gyrus | R | | 8.28 | 51 | -42 | 13 |
| Supramarginal gyrus | R | | 8.17 | 58 | -32 | 43 |
| **Superior frontal gyrus** | **R** | **21** | **9.35** | **8** | **30** | **54** |
| **Superior frontal gyrus** | **R** | **185** | **9.03** | **11** | **5** | **62** |
| Superior frontal gyrus | R | | 8.40 | 16 | 12 | 65 |
| Superior frontal gyrus | R | | 7.35 | 11 | -10 | 68 |
| Superior frontal gyrus | R | | 6.78 | 18 | -3 | 73 |
| **Insula** | **L** | **194** | **8.80** | **-44** | **10** | **-4** |
| Insula | L | | 7.86 | -46 | 2 | 4 |
| Insula | L | | 7.81 | -34 | 2 | 0 |
| Insula | L | | 6.91 | -44 | 22 | -6 |
| **Precentral gyrus** | **R** | **79** | **8.36** | **46** | **-3** | **48** |
| Middle frontal gyrus | R | | 6.85 | 43 | -3 | 57 |
| Precentral gyrus | R | | 6.53 | 51 | 2 | 40 |
| Precentral gyrus | R | | 6.49 | 33 | -8 | 48 |
| **Anterior cingulate cortex, dorsal part** | **R** | **147** | **7.97** | **11** | **17** | **35** |

| | | | | | | |
|---|---|---|---|---|---|---|
| Anterior cingulate cortex, dorsal part | L | | 7.14 | -1 | 5 | 43 |
| Anterior cingulate cortex | L | | 6.79 | -4 | 25 | 24 |
| Anterior cingulate cortex, dorsal part | R | | 6.00 | 4 | -5 | 40 |
| **Middle temporal gyrus** | **R** | **161** | **7.92** | **48** | **-60** | **13** |
| Middle temporal gyrus | R | | 7.44 | 56 | -52 | 2 |
| Middle temporal gyrus | R | | 6.88 | 56 | -57 | 10 |
| Middle temporal gyrus | R | | 6.34 | 43 | -67 | -1 |
| **Angular gyrus** | **L** | **201** | **7.84** | **-61** | **-50** | **35** |
| Supramarginal gyrus | L | | 7.80 | -64 | -47 | 26 |
| Supramarginal gyrus | L | | 7.40 | -64 | -40 | 32 |
| Central operculum | L | | 6.85 | -59 | -23 | 16 |
| **Posterior cingulate cortex** | **R** | **187** | **7.60** | **8** | **-30** | **46** |
| Precuneus | R | | 7.35 | 6 | -42 | 51 |
| Precentral gyrus | R | | 7.35 | 6 | -18 | 48 |
| Precuneus | R | | 7.15 | 8 | -57 | 62 |
| **Posterior cingulate cortex** | **L** | **24** | **7.59** | **-1** | **-25** | **26** |
| Posterior cingulate cortex | R | | 6.86 | 6 | -28 | 29 |
| **Superior occipital gyrus** | **L** | **24** | **7.39** | **-16** | **-77** | **43** |
| **Lateral occipital cortex** | **L** | **38** | **7.34** | **-31** | **-82** | **16** |
| Lateral occipital cortex | L | | 6.37 | -29 | -92 | 18 |
| **Cuneus** | **R** | **31** | **7.32** | **18** | **-82** | **26** |
| **Cuneus** | **R** | **58** | **7.30** | **13** | **-75** | **26** |
| Lateral occipital cortex | R | | 6.51 | 13 | -75 | 46 |
| Cuneus | R | | 6.23 | 13 | -75 | 35 |
| Cuneus | R | | 6.0 | 8 | -80 | 40 |
| **Lateral occipital cortex** | **R** | **61** | **7.30** | **31** | **-75** | **24** |
| **Calcarine gyrus** | **R** | **50** | **7.11** | **1** | **-70** | **13** |
| Lingual gyrus | R | | 6.44 | 4 | -80 | 2 |
| Intracalcarine cortex | L | | 6.38 | -6 | -75 | 18 |
| **Postcentral gyrus** | **L** | **20** | **6.74** | **-9** | **-47** | **57** |
| Precentral gyrus | L | | 6.33 | -14 | -37 | 43 |
| **Fusiform gyrus** | **R** | **24** | **6.67** | **36** | **-67** | **-15** |
| Fusiform gyrus | R | | 6.66 | 28 | -72 | -12 |
| **Middle frontal gyrus** | **R** | **23** | **6.51** | **31** | **40** | **26** |
| Middle frontal gyrus | R | | 6.22 | 41 | 42 | 29 |

**Result Tables for functional connectivity**

**Table S21.** Results for linear model for within and between network connectivity.

| PPI variable | FC | IV | *b* | *SE* | *t* | *p* |
|---|---|---|---|---|---|---|
| Semantic fluency > Counting | Within MDN | Intercept | 0.05 | 0.02 | 2.70 | **0.009** |
| | | Age | 0.005 | 0.03 | 0.18 | 0.85 |
| | Within DMN | Intercept | -0.002 | 0.02 | -0.07 | 0.95 |
| | | Age | 0.02 | 0.03 | 0.50 | 0.63 |
| | Between MDN and DMN | Intercept | 0.10 | 0.02 | 4.47 | **< 0.001** |
| | | Age | 0.04 | 0.03 | 1.25 | 0.22 |

Significant effects are marked in bold: $p <$ *Meff*-corrected $\alpha$ of 0.018; FC Functional connectivity; IV Independent variable; MDN Multiple-demand network; DMN Default-mode network.

**Table S22.** Results for generalized linear mixed models for within- and between-network functional connectivity effects, age, and condition on accuracy and response time.

| | | **Accuracy** | | | **Response time** | |
|---|---|---|---|---|---|---|
| *Coefficient* | *Log-Odds* | *Conf. Int (95%)* | *p* | *Estimates* | *Conf. Int (95%)* | *p* |
| Intercept | 3.09 | 2.48 – 3.70 | **< 0.001** | 6.53 | 6.48 – 6.58 | **< 0.001** |
| Within-MDN FC | -0.06 | -1.68 – 1.55 | 0.849 | -0.15 | -0.30 – 0.00 | 0.234 |
| Within-DMN FC | -0.37 | -1.93 – 1.18 | 0.621 | -0.33 | -0.50 – -0.16 | **< 0.001** |
| Between-network FC | 0.25 | -1.51 – 2.01 | 0.829 | 0.53 | 0.36 – 0.70 | **< 0.001** |
| Age | -0.01 | -0.17 – 0.16 | 0.866 | 0.07 | 0.05 – 0.08 | **< 0.001** |
| Within-MDN FC * Age | 1.55 | -1.53 – 4.63 | 0.329 | 0.98 | 0.65 – 1.31 | **< 0.001** |
| Within-DMN FC * Age | 2.10 | -0.74 – 4.93 | 0.151 | 0.90 | 0.62 – 1.19 | **< 0.001** |
| Between-network FC * Age | -2.58 | -5.59 – 0.43 | 0.096 | -0.80 | -1.11 – -0.49 | **< 0.001** |
| **Random Effects** | | | | | | |
| $\sigma^2$ | 3.29 | | | 0.13 | | |

| | | |
|---|---|---|
| $\tau_{00}$ | 0.22 $_{Subj}$ | 0.01 $_{Subj}$ |
| | 1.71 $_{Category}$ | 0.00 $_{Category}$ |
| ICC | 0.37 | 0.12 |
| N | 58 $_{Subj}$ | 52 $_{Subj}$ |
| | 20 $_{Category}$ | 20 $_{Category}$ |
| Observations | 9837 | 9675 |
| Marginal $R^2$ / Conditional $R^2$ | 0.002 / 0.371 | 0.027 / 0.143 |

Significant effects are marked in bold. Contrasts are sum coded. P-values were obtained via likelihood ratio tests. Conf. Int. Confidence interval.

**Table S23.** Results of post-hoc tests for two-way interactions Age x Connectivity measure for response time model. P-values are Bonferroni-corrected.

| **Contrast** | **FC** | ***Estimate*** | ***SE*** | ***df*** | ***Conf. Int (95%)*** | ***z*** | ***p*** |
|---|---|---|---|---|---|---|---|
| OA – YA | Within MDN | 661 | 116 | Inf | 434 – 887 | 5.72 | **< 0.001** |
| | Within DMN | 602 | 100 | Inf | 405 – 799 | 6 | **< 0.001** |
| | Between MDN and DMN | -524 | 110 | Inf | -739 – -308 | -4.76 | **< 0.001** |

Significant effects are marked in bold. FC functional connectivity; SE standard error; df degrees of freedom; Conf. Int confidence intervals.
